# Packaged delivery of CRISPR-Cas9 ribonucleoproteins accelerates genome editing

**DOI:** 10.1101/2024.10.18.619117

**Authors:** Hannah Karp, Madeline Zoltek, Kevin Wasko, Angel Luis Vazquez, Jinna Brim, Wayne Ngo, Alanna Schepartz, Jennifer Doudna

## Abstract

Effective genome editing requires a sufficient dose of CRISPR-Cas9 ribonucleoproteins (RNPs) to enter the target cell while minimizing immune responses, off-target editing and cytotoxicity. Clinical use of Cas9 RNPs currently entails electroporation into cells *ex vivo*, but no systematic comparison of this method to packaged RNP delivery has been made. Here we compared two delivery strategies, electroporation and enveloped delivery vehicles (EDVs), to investigate the Cas9 dosage requirements for genome editing. Using fluorescence correlation spectroscopy (FCS), we determined that >1300 Cas9 RNPs per nucleus are typically required for productive genome editing. EDV-mediated editing was >30-fold more efficient than electroporation, and editing occurs at least two-fold faster for EDV delivery at comparable total Cas9 RNP doses. We hypothesize that differences in efficacy between these methods result in part from the increased duration of RNP nuclear residence resulting from EDV delivery. Our results directly compare RNP delivery strategies, showing that packaged delivery could dramatically reduce the amount of CRISPR-Cas9 RNPs required for experimental or clinical genome editing.

**Graphical Abstract:** 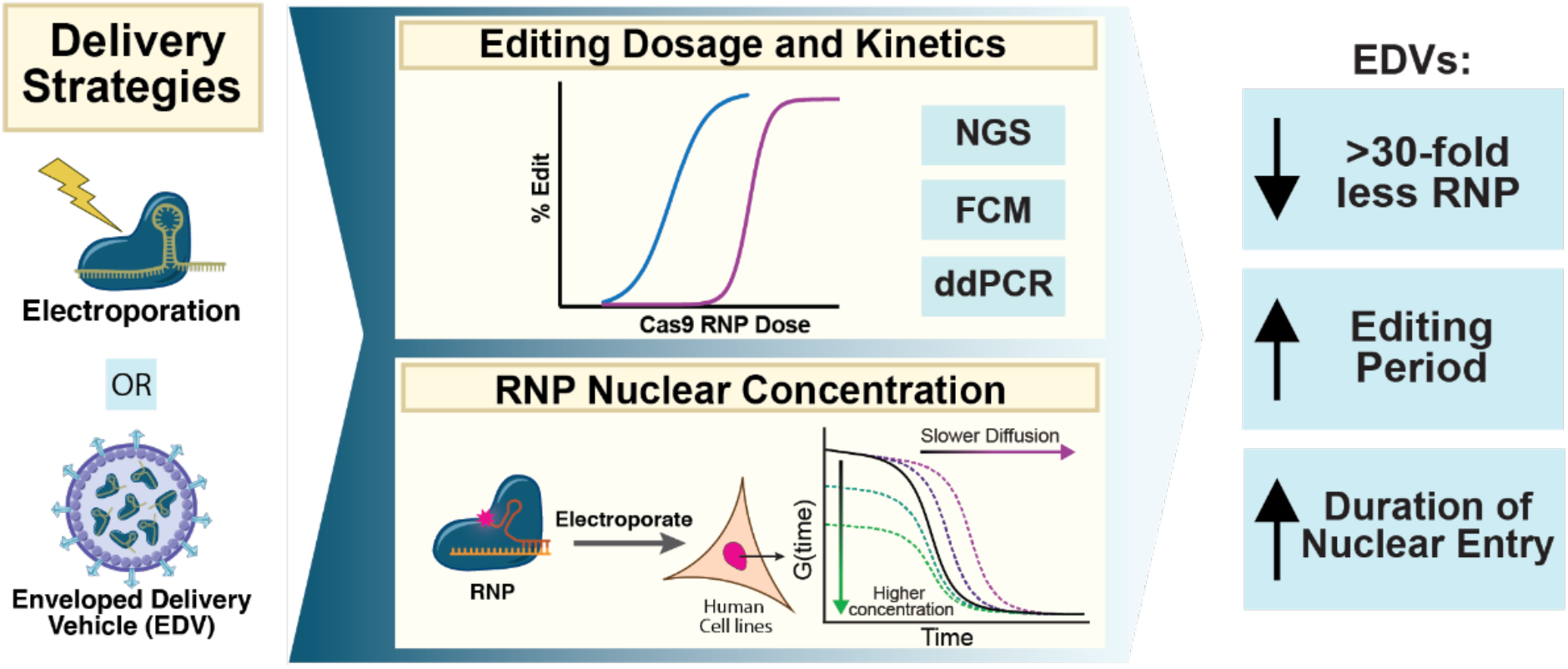

## Introduction

CRISPR-based genome editing therapies have enormous potential to cure genetic diseases. Despite this promise, safe and effective delivery of genome editors remains a challenge for both therapeutic development and fundamental research (1, 2). Broadly speaking, genome editors can be delivered either as a nucleic acid, to be transcribed and/or translated in the target cell, or as an intact ribonucleoprotein complex (RNP) (1). There are distinct advantages to delivering genome editors as RNPs, including shorter intracellular lifetimes that minimize off-target edits (3) and reduce immunogenicity (4–6). Compared to mRNA delivery, RNP delivery may result in lower levels of toll-like receptor activation (4, 7), and enable higher *in vivo* editing efficacy by bypassing *in situ* translation of mRNA (8) and protecting the single guide RNA (sgRNA) integrity due to Cas9 protein binding (9). RNP delivery also avoids risks of random DNA integration posed by viral vectors, including lentivirus and adeno-associated virus (AAV) (10, 11).

While delivery of proteins and RNA to the interior of cells remains a critical therapeutic challenge (12), extensive engineering efforts have generated multiple promising *ex vivo* and *in vivo* Cas9 RNP delivery strategies (1, 2). RNP electroporation is the most common strategy, with widespread use in CRISPR genome editing therapies (13, 14). RNP electroporation is less cytotoxic than nucleic acid electroporation (15), is efficient in primary cells and has higher specificity than delivery systems that result in extended genome editor expression (3). However, RNP electroporation requires an *ex vivo* approach, limiting its therapeutic utility. Furthermore, electroporation can impact cell viability (16) and lead to high manufacturing costs for cell-based therapies (16). Alternatively, enveloped delivery vehicles (EDVs) derived from human immunodeficiency virus (HIV) offer a packaged approach to Cas9 RNP delivery (17). EDVs leverage HIV’s intrinsic intracellular delivery capabilities and production scalability while mitigating the risk of lentiviral genome integration or extended transgene expression (17–19). EDVs use vesicular stomatitis virus glycoprotein, VSVG, for cellular uptake and endosomal escape, which typically exhibits broad cell tropism (11, 17, 18). Recent work demonstrated that binding-deficient VSVG combined with antibody-derived targeting motifs enables cell-type specific Cas9 delivery both *ex vivo* and *in vivo* (18, 20). However, despite the promise of RNPs and EDVs, little is known about how much functional Cas9 RNP can be delivered in each case and how much is required for efficient editing in human cells (3, 6, 16, 21).

To address these questions, we compared electroporation and EDVs for the delivery of S. *pyogenes* Cas9 RNPs to edit various human cell types. We determined the impact of delivery modality on the rate of DNA cleavage and repair. Using fluorescence correlation spectroscopy (FCS), we found that >1300 Cas9 RNPs per nucleus are required for editing in human cell lines. At comparable Cas9 RNP doses, EDVs are 30-to 50-fold more effective at editing, across multiple human cell types and target genome sequences. Furthermore, EDV delivery generates genome edits twice as fast as electroporation. We hypothesize that the observed differences in editing efficacy and rate result in part from differences in RNP versus EDV trafficking to the cell nucleus. Our results suggest that the Cas9 RNP dosage used for current *ex vivo* research and clinical genome editing could be substantially reduced by switching from electroporation to a packaged delivery strategy such as EDVs. These findings also reveal the importance of delivery modality for genome editing efficacy and pave the way for engineering optimal delivery methods to ensure maximal genome editing with minimal side effects.

## Results

### Several thousand Cas9 RNPs per cell nucleus are sufficient for editing human cell lines

To estimate the number of Cas9 RNP molecules per cell nucleus that are sufficient for editing, we used fluorescence correlation spectroscopy (FCS) to measure both the concentration and rate of diffusion of fluorescently labeled RNPs in cells (Fig. 1A) (22–24). We incubated purified Cas9 protein with a dual guide RNA consisting of commercially available ATTO^TM^ 550-labeled tracrRNA and a *B2M*-targeting crRNA (Supplemental Table 1). As we were most interested in quantifying the amount of intact RNP, we reasoned that fluorescently labeled guide RNA would provide a detectable shift in diffusion time between free RNA and the intact RNP in buffer (Fig. S1A, B) and in cells. We compared the concentration and diffusion rates of electroporated Cas9 pre-complexed with the labeled dual-guide RNA versus the labeled dual-guide RNA alone in HeLa cells (Fig. 1A). After 24 hours, fluorescent signal in nuclei was evaluated by FCS, with observed RNP diffusion time significantly higher than for dual-guide RNA alone (2.0 vs 1.0 ms, p-value =0.0004, Fig. 1B). The nuclear concentration of the RNP condition was higher than that observed for the dual-guide RNA only condition (7.5×10^7^ Cas9 per cell, 25 vs. 19 nM, Fig. S1C), possibly due to both nuclear localization and RNA stabilization that results from Cas9 binding (25). Two-component analyses of nuclear autocorrelation functions obtained by FCS provided best fits to the data and were used for all future nuclear delivery analyses (Fig. S1D, E).

**Figure 1:**
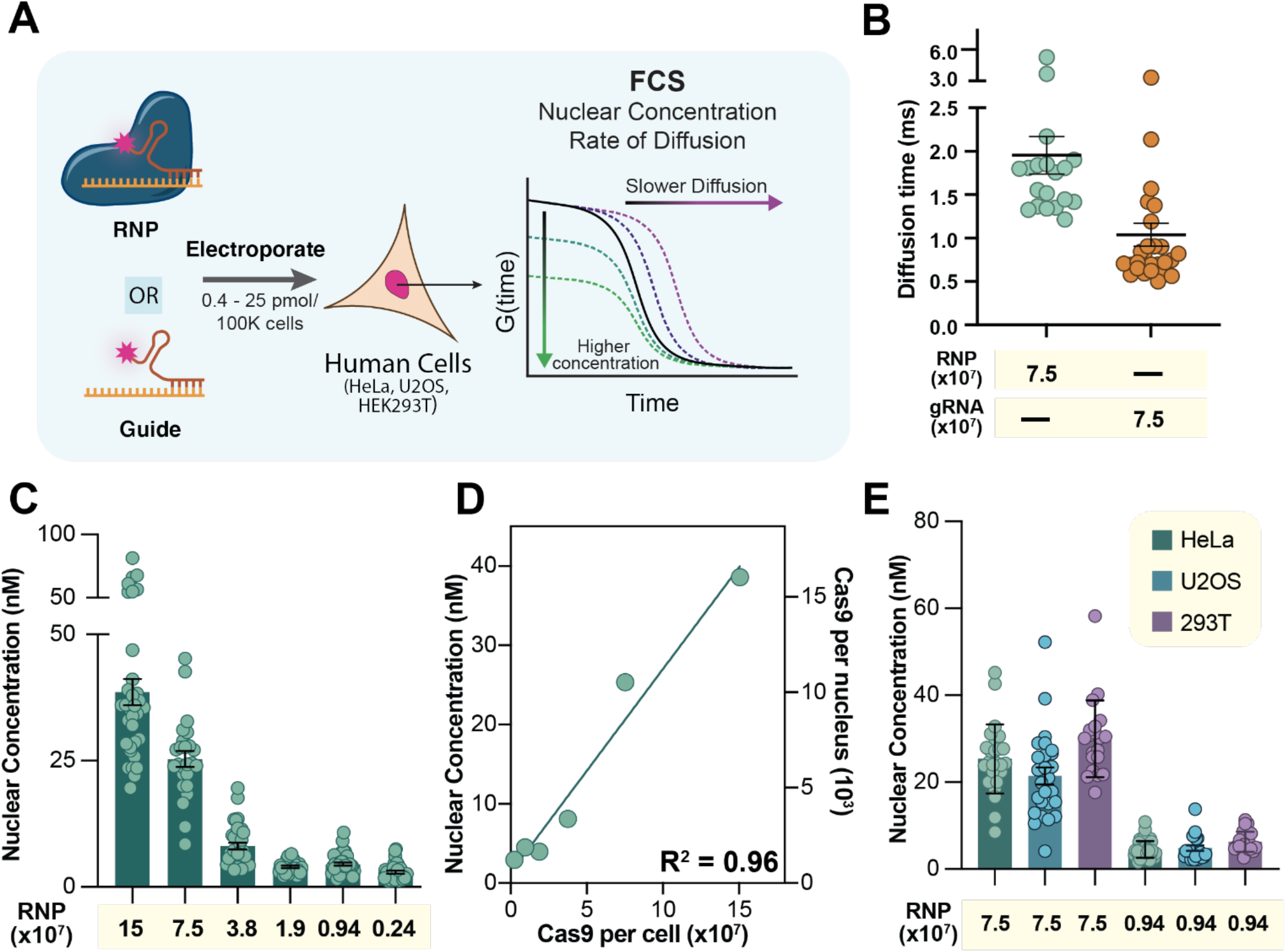
Quantifying Cas9 RNP nuclear concentration delivered by electroporation with fluorescence correlation spectroscopy (FCS) A) Experimental schematic of workflow to quantify the Cas9 RNP nuclear concentration required for editing. B) Diffusion time of Cas9 RNP or gRNA, in Cas9 per cell, delivered in Hela cells and measured at 24 hrs (2.0 vs 1.0 ms, p-value =0.0004) Each point represents the average diffusion time in an individual cell modeled with a two-component diffusion fitting (Fig. S1). FCS diffusion times provided in ms, n>25 for each FCS condition with at least two biological replicates each (mean ± SEM). C) FCS analysis of HeLa cells electroporated with the Cas9 RNP. Nuclear concentration of Cas9 RNP as a function of dosage (in Cas9 per cell). Each point represents the concentration in an individual cell. FCS values provided in nM, n>25 for each FCS condition with at least two biological replicates each (mean ± SEM). All concentration values and diffusion times were derived by fitting FCS traces with a two-component 3D diffusion equation (see Methods for more detail). D) (Left Axis) Average nuclear concentration of Cas9 RNP versus dosage shows a strong linear correlation. (R^2^ = 0.96). (Right Axis) Estimated number of Cas9 per nucleus from nuclear concentration values calculated by FCS and volume of HeLa nucleus (690 μM^3^) (46). E) FCS analysis of nuclear concentration for HeLa, U2OS, and HEK293T cells. FCS values provided in nM, n>20 for each FCS condition with at least two biological replicates each (mean ± SEM). Exact values for FCS, including experimental and biological replicates, mean, and SEM are reported in Supplemental Table 2.

Next, we quantified the nuclear concentration of Cas9 RNA containing a *B2M*-targeting, fluorescently labeled dual-guide RNA, across a range of dosages (15×10^7^ to 0.24×10^7^ Cas9 per cell) in HeLa cells at 24 hrs (Fig. 1C, D). The RNP nuclear concentrations resulting from electroporation were linear across this dose range (R^2^=0.96) ranging from 39 nM to 3 nM (Fig. 1C). Using published estimates of HeLa nuclear volume, ∼690 μM^3^ (34), we calculated the number of Cas9 molecules per nucleus to be 16,000-1200 molecules for this dose range (Fig. 1C, D; Supplemental Table 2).

We wondered whether electroporation delivery efficiency varies substantially by cell type. Measurement of Cas9 RNP nuclear concentrations in two additional cell lines, HEK293T and U2OS, at two doses (7.5×10^7^ and 0.94×10^7^ Cas9 per cell) revealed similar values to those measured in HeLa cells (Fig. 1E). This suggests that Cas9 RNP delivery efficiency by electroporation in multiple cell types is similar and thus the large RNP dose difference required for editing in different cell types may be due to differences in epigenetic landscape or DNA repair mechanisms (26, 27).

### EDVs dramatically reduce the amount of Cas9 RNP needed for genome editing

To determine how much functional Cas9 RNP is required for efficient editing in human cells, we first determined how much total Cas9 protein is needed for editing when delivered by electroporation or EDV into HEK293T or HeLa cells (Fig. 2A; Fig. S2). For electroporation, the dosage that resulted in half maximal editing of the B2M gene (EC50) was 2.4×10^6^ and 7.3×10^6^l Cas9 per cell in HEK293T and HeLa cells, respectively (Fig. 2B). For EDV delivery, the EC50 was 5.7×10^4^ and 1.5×10^5^ Cas9 per cell in HEK293T and HeLa cells, respectively (Fig. 2B). These values correspond to 42- and 50-fold reductions in required Cas9 dose for HEK293T and HeLa cells, respectively, for EDV-mediated RNP delivery compared to electroporation.

**Figure 2:**
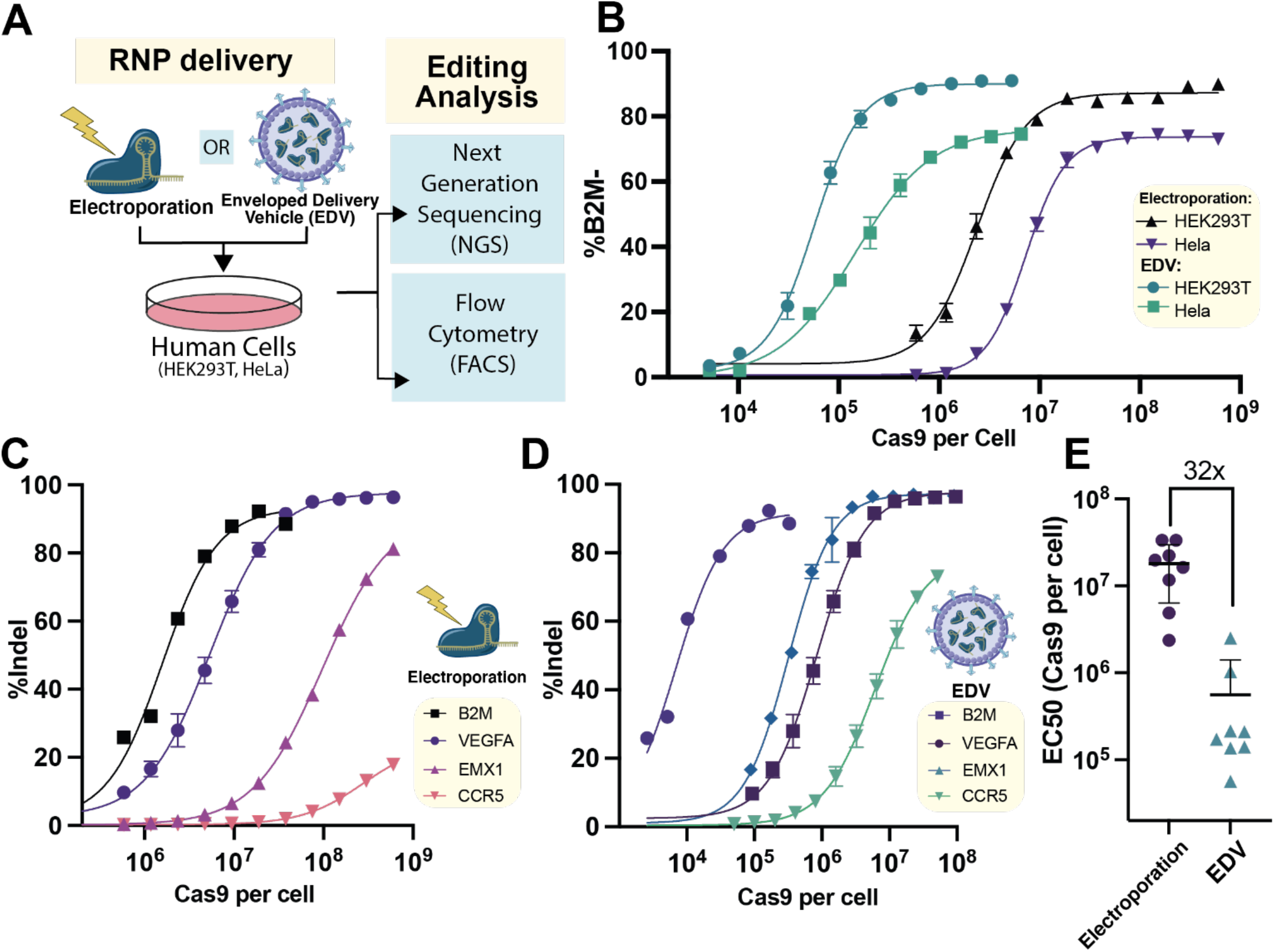
Assessing dosage requirements of Cas9 RNP delivered by electroporation and EDVs. **(A)** Experimental schematic of workflow to quantify the Cas9 RNP doses required for editing by electroporation and EDVs. **(B)** To assess Cas9 RNP dosage required for editing in HEK293T, Hela, Jurkats, and T cells were treated with varying doses of B2M-targeting Cas9 by electroporation and EDVs. Analysis was performed by flow cytometry 4 days post treatment to assess B2M knockdown. **(C)** Electroporated and **(D)** EDV delivery of Cas9 RNP targeting the B2M, VEGFA, EMX1 and CCR5 loci in HEK293T cells. Analysis was performed by next generation sequencing (NGS) 4 days post treatment to assess indels. **(E)** Comparison of required RNP doses delivered by electroporation and EDV for 9 different B2M guides in HEK293T (p-value = 0.0003, Mann Whitney Test). Analysis was performed by flow cytometry 4 days post treatment to assess B2M KO. n=3 technical replicates were used in all experiments. Datapoints represent the mean with error bars displaying SD. RNP dose curves were modeled (Prism v10) as sigmoidal (4PL, X is concentration).

Extensive work has shown that the choice of sgRNA strongly impacts the activity and specificity of Cas9 (28, 29), but it remains unknown the degree to which the sgRNA impacts the Cas9 RNP dosage required for editing. We compared doses required for editing using four different guide RNAs targeting *B2M, VEGFA*, CCR*5* and *EMX1* and found that the amount of Cas9 required for editing varied by greater than 100-fold depending on guide choice (Fig. 2C, 2D).

We wondered whether these differences in required RNP dosage by these sgRNAs were due to differences between the *B2M, VEGFA, CCR5* and *EMX1* loci. Therefore, to minimize the impact of potential differences in chromatin state, we generated a panel of nine sgRNAs targeting a ∼200-bp window in the *B2M* locus and compared the dosages required for *B2M* knockout in HEK293T and HeLa cells (Fig. 2E; Fig. S3). Different sgRNAs resulted in substantial differences (>100 fold) between RNP doses required for editing using either electroporation or EDVs. For electroporation and EDVs respectively, the EC50 values across sgRNAs were highly correlated between cell lines (R^2^ = 0.78, R^2^ = 0.83) (Fig. S4A, B). Interestingly however, the sgRNA trends were only loosely correlated across delivery modalities (R^2^ = 0.55, Fig. S4C) and there was no correlation between individual sgRNAs’ maximum editing levels and dosage requirements (R^2^ =0.01, R^2^ =0.08, Fig. S4D, E).

To remain functional in human cells Cas9 must remain guide-complexed and retain biochemical cleavage activity. Typically, ∼20-40% of purified Cas9 is active *in vitro* (30–32). Across three sgRNAs, our in-house purified Cas9 averaged 30% activity (Fig. S5), but we wondered whether EDV packaging impacts the activity of encapsulated Cas9 RNPs. First, we quantified the maximum percent of Cas9 protein in EDVs that could be complexed with sgRNA by measuring the Cas9 protein and sgRNA concentrations using Cas9 ELISA and RT-qPCR, respectively (Fig. S6A). Across five independent batches of EDVs, only 8.3 +/-1.7 sgRNA molecules were measured for every 100 molecules of Cas9 protein (Fig. S6A).

We first tested whether sgRNA availability limits the Cas9 cleavage functionality in EDVs. Cas9 packaged in EDVs with or without sgRNA was introduced into cells that were also transfected with a plasmid expressing a *B2M*-targeting sgRNA (Fig. S6B). However, this additional guide RNA supplementation failed to result in measurable improvement in editing efficacy (Fig. S6B). We next tested whether the Cas9 protein in EDVs is degraded or unfolded, which could render it incapable of guide RNA binding. Using Western blotting, we determined that most Cas9 in EDVs remains uncleaved from the lentiviral polyprotein Gag or is degraded in EDVs, and that only 35% of the Cas9 is intact (Fig. S6C, D). Together with previous results showing that Cas9 protein alone readily denatures at 37°C (33), this finding suggests that the vast majority of Cas9 in EDVs is not functional. Furthermore, we conclude that the per-molecule difference in RNP delivery efficiency between EDVs and electroporation is substantially greater than the >30-fold difference measured in editing assays (Fig. 2).

### EDV-mediated Cas9 RNP delivery results in rapid DNA cleavage and repair

The efficiency of genome editing depends on the integrated amount and duration of functional Cas9 RNP residing in the cell nucleus. To gain insight into the rate and duration of Cas9 RNP activity as a function of delivery modality, we treated cells with a range of saturating dosages (defined as that required for 90% maximal editing) of *B2M*-targeting RNP delivered by either electroporation or EDVs (Fig. 3A). Cells were harvested over the course of 54 hours post-treatment and analyzed using digital droplet PCR (ddPCR) and next generation sequencing (NGS) to measure double-strand DNA breaks (DSBs) and genome edits, respectively (Fig. 3A, B). Electroporation permeabilizes mammalian cells using electric pulses lasting a few microseconds to induce pore formation in the plasma membrane. These pores reseal within seconds to minutes, however, limiting the time in which RNPs can access the cell interior (34). Consistent with this transient delivery window, at all RNP amounts tested, electroporation resulted in the majority of observed DSBs occurring within two hours post delivery (Fig. 3C). The highest and lowest (6×10^7^ and 0.75×10^7^ Cas9 per cell) RNP doses resulted in similar maximum concurrent DSBs (64% versus 54%, respectively) (Fig. 3C).

**Figure 3:**
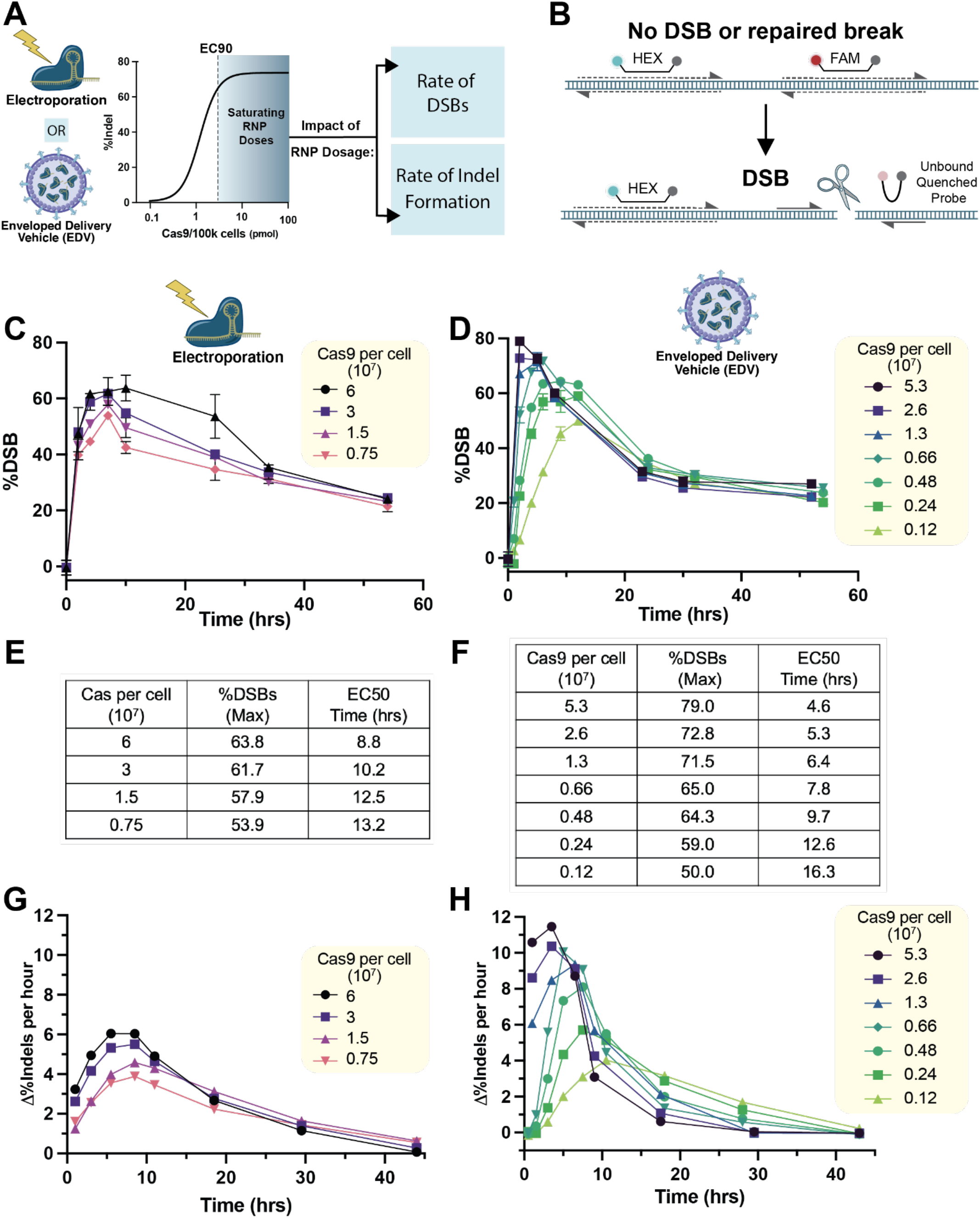
Kinetics of double stranded breaks and DNA repair resulting from delivery by RNP electroporation and EDVs. **(A)** Schematic overview of time course experiment comparing the impact of saturating, defined as doses at or exceeding levels for 90% of the maximum editing (>EC90), doses of Cas9 RNP delivered by electroporation and EDVs on the rate of double stranded breaks (DSBs) and consequent indel repair detected by NGS. **(B)** Experimental setup of the digital droplet PCR used to quantitatively detect DSBs. A HEX probe spans the first amplicon which is centromere proximal to the break site. A second FAM probe anneals to the second amplicon, which is lost upon DSB or chromosome loss. DSBs caused by **(C)** electroporation and **(D)** EDV delivery of Cas9 RNP targeting the B2M locus over a 54 hour time frame. Tables comparing the maximum percentage of synchronous DSBs and timeframe for half maximal editing (EC50) resulting from delivery of *B2M*-targeting RNP in HEK293T by **(E)** electroporation and **(F)** EDV delivery. Rate of indel formation caused by **(G)** electroporation and **(H)** EDV delivery of *B2M*-targeting RNP in HEK293T cells measured by NGS (Raw NGS time courses in Supplemental figure 7). All doses are at levels that meet or exceed the amount necessary for 90% maximal editing. n=3 technical replicates were used in all experiments. Datapoints represent the mean with error bars displaying SD.

We initially hypothesized that EDVs delivery would take longer than electroporation because EDV endocytosis requires fusion with endosomes to deliver RNPs into the cytosol. Interestingly, at a high RNP dose (5.3×10^7^ Cas9 per cell), EDV-mediated RNP delivery occurred as quickly as electroporation, resulting in most DSBs occurring within two hours (Fig. 3D). These data show that the minimal time frame for EDV-mediated Cas9 RNP intracellular delivery, nuclear localization and genome target cleavage occurs within two hours, despite requiring additional delivery steps. Notably, for EDVs the timeframe for DSB formation was substantially more dose dependent, with the lowest doses of Cas9 delivered by EDV requiring 12 hours to reach the maximum level of DSBs (49%) (Fig. 3D). Importantly, this shows that it is possible to control and tune the rate of editing with the concentration of EDV.

In the absence of a DNA donor template, DSBs are typically resolved through non-homologous end joining (NHEJ). It remains unclear whether the time frame for DNA repair is impacted by delivery mode, particularly because delivery can perturb cellular metabolism and viability (16, 34). At high RNP doses (6×10^7^ and 5.3×10^7^ Cas9 per cell), both strategies result in similar timeframes for DSB formation, but EDV delivery generated genome edits twice as fast (Fig. 3E-H; Fig. S7). At these same doses, half maximal editing for electroporation and EDVs occurred by 8.8 and 4.6 hours, respectively (Fig. 3E, F; Fig. S7). For electroporation, genome editing occurred within a similar timeframe for all doses, with the maximum rate of editing occurring in the 6-8 hour window (Fig. 3G). Conversely, the rate of editing for EDVs was highly dose dependent (Fig. 3H). At the earliest time point (one hour), the rate of genome edits was three times higher for EDVs than for electroporation at comparable doses, 10.6 vs 3.2 %edits/hour, respectively (Fig. 3G, H; Fig. S7). At lower EDV doses, the rate of editing remained high until 28 hours following EDV treatment (Fig. 3H), consistent with the extended duration of DSB formation (Fig. 3D). Combined, these results suggest that EDVs can continue to deliver functional RNPs into the nucleus over a prolonged time period.

### EDV delivery results in extended nuclear accumulation of Cas9 RNPs over time

We hypothesized that the difference in genome editing kinetics observed using electroporation versus EDV-mediated Cas9 RNP delivery may correspond to the length of time that Cas9 RNPs reside in the nucleus. We hypothesized that EDVs could continue to deliver Cas9 RNPs to the nucleus over an extended time frame, whereas electroporation would result in decreasing levels of nuclear Cas9 following initial delivery. To quantify electroporated Cas9 RNPs as a function of time, we used FCS to measure the nuclear concentration of Cas9 RNP in HeLa cells at 12, 24 and 36 hours following electroporation. At two different electroporation doses (7.5×10^7^ and 0.94×10^7^ Cas9 per cell), nuclear RNP concentrations did not vary at the three timepoints (Fig. 4A; Supplemental Table 2). We used confocal microscopy to visualize the electroporated Cas9 RNPs in HeLa cells at these time points and validated the results by quantifying relative nuclear concentrations of Cas9 using fixed confocal microscopy (Fig. 4B; Fig. S8A, see Methods). Consistent with our editing time course (Fig. 3), the nuclear concentration of Cas9 delivered by EDVs continued to increase for up to 32 hours after delivery (Fig. 4C, D). Interestingly, for EDV delivery, Cas9 appeared to also accumulate around the nuclear envelope which was not observed for electroporation (Fig. 4B, Supplemental Table 3). This may be due to the nuclear export signals present on the uncleaved Gag-Cas9 construct that facilitates EDV packaging (Fig. S6D, S9) (18) causing some of the Cas9 to accumulate around the nuclear envelope.

**Figure 4:**
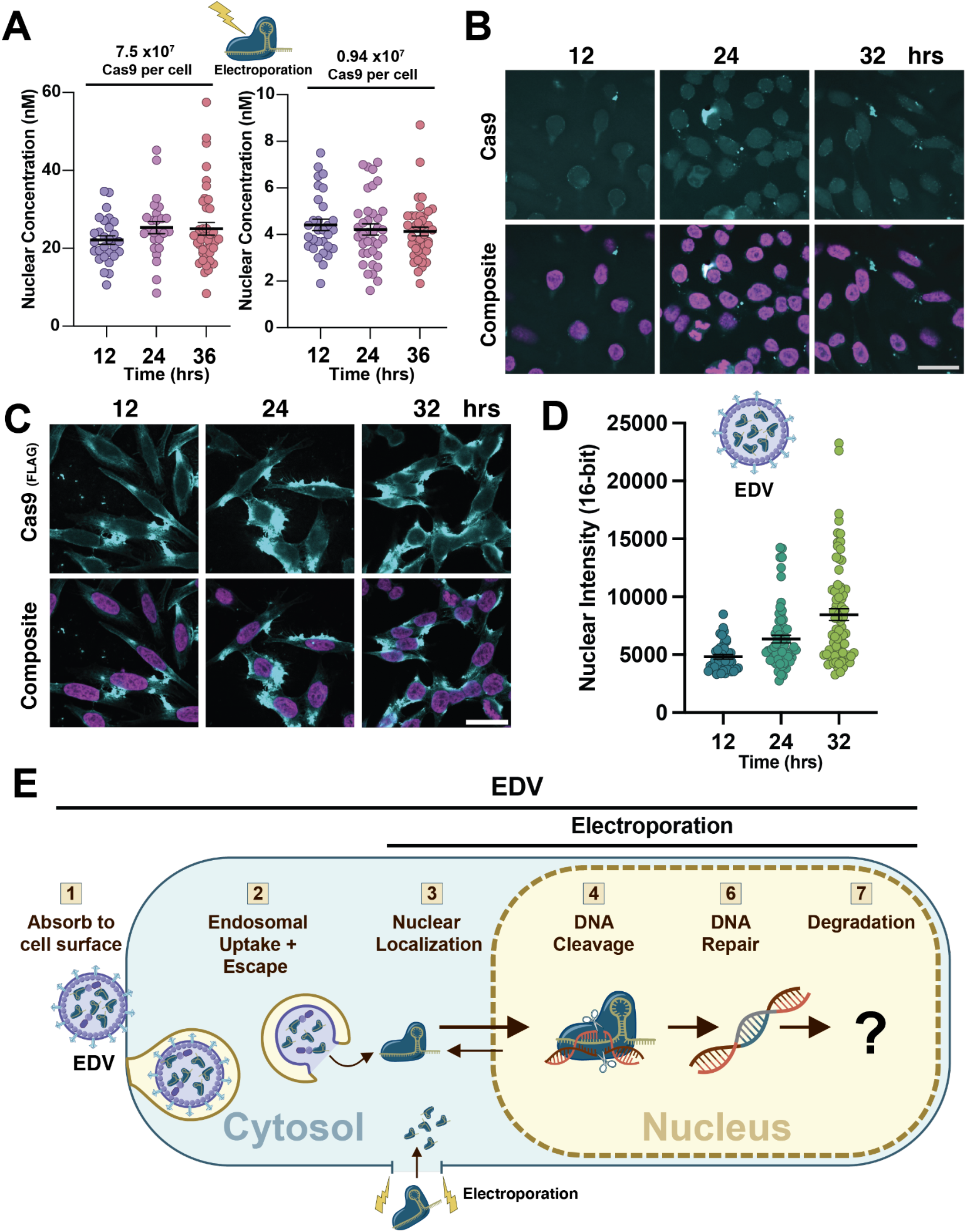
Comparison of nuclear localization and concentration of Cas9 delivered by electroporation and EDVs as function of time. **(A)** Quantification of the nuclear concentration of Cas9 RNP in HeLa cells by fluorescence correlation spectroscopy (FCS) at 12, 24, and 36 hours post-electroporation at two doses, 7.5×10^7^ and 0.94×10^7^ Cas9 per cell. Each point represents the concentration in an individual cell. FCS values provided in nM, n>25 for each FCS condition with at least two biological replicates each (mean ± SEM). All concentration values and diffusion times were derived by fitting FCS traces with a two-component 3D diffusion equation (see Methods). Representative fixed confocal images of HeLa cells stained for Cas9 (top) and composite image of Cas9 overlaid with nuclear counterstain showing the nuclear intensity of Cas9 at 12, 24 and 36 hours following **(B)** electroporation (1.2×10^8^ Cas9 per cell) and **(C)** EDV treatment (∼3.0×10^6^ Cas9 per cell). Scale bar is 25 uM. **(D)** Relative quantification of median nuclear intensity of Cas9 delivered by EDV at 12, 24 and 36 hours (see Methods). Intensity on 16-bit scale (0 to 65536). Each data point represents an individual nuclei n> 40 (mean ± SEM). **(E)** Schematic overview of Cas9 delivery by EDVs and Electroporation.

Collectively, these results together with insights from previous studies support a model for how EDV delivery impacts the rate and mode of Cas9 RNP delivery (Fig. 4E). The earliest detectable genome editing occurred within two hours for all EDV doses studied (Fig. 3), demonstrating that Cas9 RNP nuclear entry occurs quickly following delivery. EDVs adsorb quickly to the cell membrane (35) and remain associated even after cell washing. Over time, these EDVs undergo endosomal uptake and escape mediated by VSV-G on the EDV surface (35). Cas9 RNP encapsulation by EDVs appears to extend RNP half-life in the nucleus, possibly by impacting Cas9 degradation mechanisms.

## Discussion

Safe and effective delivery of CRISPR-Cas9 based genome editing enzymes has profound potential to advance both therapeutic development and fundamental research. Focusing on Cas9 RNP delivery, we compared electroporation and enveloped delivery vehicles (EDVs) for their ability to introduce sufficient RNPs into cells to mediate intended genome modifications. Our data show that a minimum of ∼1300 Cas9 molecules per cell nucleus are required for half maximal genome editing across a range of different human cell lines. We also found that increasing genome editor lifetime in the nucleus is critical to minimize effective RNP concentration while maintaining editing efficacy. Recent work demonstrated that inhibiting Cas9 Keap1-mediated degradation enhanced epigenome editor performance (36). Future research should investigate whether thermostable Cas9 variants, such as iGeoCas9 (33), exhibit increased nuclear half-life, or for EDV delivery, allow for a larger proportion of RNPs to be functional.

We found that packaged delivery of Cas9 RNPs within EDVs resulted in continued RNP nuclear localization over a prolonged period (Fig. 4C, D). Similarly, other RNP delivery strategies, including lipid nanoparticles and cell-penetrating peptides, may result in prolonged delivery windows compared to direct RNP electroporation. Importantly, these extended time windows could be leveraged for spatiotemporal control (37) or to impact DNA repair outcomes (16, 37). Furthermore, the delay between DSB and subsequent genome editing will likely be cell-type dependent, with longer repair times expected for clinically relevant post-mitotic cell types such as neurons and cardiomyocytes (37).

Improving nuclear localization efficiency of the Cas9 RNP is of paramount importance. Our study indicates that EDV delivery protects the Cas9 RNP and increases its delivery duration. For EDV delivery, there was noticeable accumulation of Cas9 around the nuclear envelope which suggests that nuclear localization may be limiting. This is consistent with recent EDV engineering demonstrating that adding additional nuclear localization signals (NLSs) to Cas9 can improve EDV-mediated editing efficiency by ∼2-fold (38). For RNP electroporation, previous studies have also illustrated the importance of NLS optimization (16, 38), but how NLS tiling impacts RNP activity, cellular perturbations or half-life, and whether the optimal NLS configuration is different for genome editor fusion constructs (i.e. base or prime editors), or other Type II and Type V Cas nucleases, remains to be tested.

We show that understanding the impact of delivery modality on RNP intracellular trafficking, localization and genome editing efficacy can identify delivery bottlenecks that could be the focus of future engineering of improved RNP-based gene therapies. Our study examined the dosage requirements of nuclease active SpyCas9 in human cell lines. Future work should investigate the editor dosage requirements for other clinically-relevant cell types, as well as whether or how these requirements differ for other editing tools used for base, prime, and epigenome editing, to best preserve genomic and cellular integrity while efficiently achieving the desired genomic alterations.

## Materials and Methods

### Plasmid construction

Restriction enzymes used in this study were purchased from New England Biolabs (NEB). Plasmids were constructed using NEBuilder HiFi DNA Assembly Master Mix (NEB) with PCR products and backbone restriction digests. For guide plasmid cloning, protospacer oligos were annealed and then inserted using BsmBI golden gate assembly into the optimized Gag-Cas9 (Gag-3xNES-2xNLS-Cas9-U6-sgRNA) and PsPax-U6-sgRNA plasmids as previously reported (18). Oligos encoding the sgRNA spacers (IDT), and all other oligos used in this study, are outlined in Supplementary table 3.

Cloning and DNA preparations were performed in MultiShot StripWell Mach1 (ThermoFisher). All plasmids used in tissue culture were prepared with Qiagen Plasmid Maxi Kit (Qiagen). All plasmids were sequence confirmed prior to use (Plasmidsaurus, UC Berkeley DNA Sequencing Facility).

### Tissue culture

Lenti-X (Takara Biosciences), HEK293T, U2OS, and HeLa cells were obtained and authenticated by the UC Berkeley Cell Culture facility. Lenti-X, HEK293T, U2OS and HeLa cell lines were cultured in DMEM (Fisher Scientific) supplemented with 100 U/ml Penicillin-Streptomycin (Thermofisher), and 10% (v/v) fetal bovine serum (FBS) and passaged with Trypsin-EDTA (0.25%, Phenol Red, Fisher Scientific).

### RNP electroporation

sgRNA (IDT, Supplemental Table 1) were resuspended in IDT duplex buffer to 100 μM concentration. Cas9 RNPs were formed by combining the sgRNA and 40 μM Cas9-NLS (UC Berkeley QB3 MacroLab) at a molar ratio of 1.5:1 and incubating at room temperature for 10-15 min. Electroporation was performed using a 96-well format 4D-nucleofector (Lonza) with 10^5^ cells per well (unless otherwise specified). HEK293T cells were electroporated with the SF buffer and the CM-130 pulse code. HeLa cells were electroporated with SE buffer and the CN-114 pulse code. For cell lines, cells were immediately resuspended in pre-warmed media and transferred to culture plates.

### EDV and Lentiviral production

Cas9-EDVs were produced as previously described (17, 18). Briefly, Cas9-EDVs were produced by seeding approximately 4 million Lenti-X cells (Takara Bio) into 10 cm tissue culture dishes (Corning) and transfecting the next day with 1 μg pCMV-VSV-G (Addgene plasmid #8454), 6.7 μg Gag-Cas9-U6-sgRNA, 3.3 μg psPax2-U6-sgRNA (Addgene plasmid #12260) using TransIT-LT1 (Mirus Bio) at a 3:1 TransIT-LT1:plasmid ratio. Two days post-transfection, Cas9-EDV-containing supernatants were harvested, passed through a 0.45 μm PES syringe filter (VWR) and concentrated with ultracentrifugation by laying EDV containing supernatant on top of 30% sucrose in 100 mM NaCl, 10 mM Tris-HCl (ph 7.5), 1 mM EDTA at 25,000 rpm with a SW28 rotor (Beckman Coulter) for 2 hrs at 4°C in polypropylene tubes (Beckman Coulter). Concentrated Cas9-EDVs were resuspended in Opti-MEM (Gibco) at a final concentration of 20x unless otherwise noted and frozen at -80 °C until use.

### Fluorescence correlation spectroscopy

On the day of each FCS experiment, 300 uL of 10 – 100 nM AlexaFluor 594 hydrazide for electroporation experiments using ATTO^TM^ 550 for Cas9 detection, both diluted into MilliQ, was added to one well of the 8-well microscopy dish and incubated at 37°C for at least 30 min. Immediately prior to measurements, a DNA stain, 300 nM Hoechst 33342, was incubated with the cells for 5 minutes to visualize nuclei. After nuclear dye incubation, cells were washed 2x with DPBS and incubated with pre-warmed DMEM for imaging.

The general procedures used for FCS have been described previously (39–41). Experiments were performed with a STELLARIS 8 microscope (Leica Microsystems) with a Leica DMi8 CS scanhead, a HC Plan-Apo 63x/1.4NA water immersion objective, and a pulsed white-light laser (440 nm-790 nm; 440 nm: > 1.1 mW; 488 nm: > 1.6 mW; 560 nm: > 2.0 mW; 630 nm: > 2.6 mW; 790 nm: > 3.5 mW, 78 MHz). All confocal imaging was performed using HyD S or HyD X detectors in counting mode, while FCS measurements were carried out using only a Hybrid HyD X detector in counting mode. All microscopy experiments were performed at 37°C (monitored using Oko-Touch) and 5% CO_2_ in a blacked out cage enclosure (Okolab). Before each experiment, the correction collar of the objective was adjusted by maximizing the counts per molecule for the AlexaFluor 594 hydrazide dye standard; minor fluctuations in the correction collar are expected based on the variable thickness of the glass-bottom microscopy dishes (LabTek^TM^). After correction collar adjustment, ten five-second autocorrelation traces were obtained from the well containing dye standard to calculate the focal volume of the microscope (see Analysis of FCS Data for more detail). AlexaFluor 594 was measured using the same settings as ATTO^TM^ 550.

For electroporation experiments, ATTO^TM^ 550 was excited at 553 nm with an emission window of 570 - 660 nm, and Hoescht 33342 was excited at 405 nm with an emission window of 432 – 509 nm. Laser intensity for ATTO^TM^ 550 was determined using *in vitro* samples of each respective protein and guide and determining the maximal laser intensity for which the observed counts per molecule (CPM) remained within a linear range. The pinhole S5 of the laser was set to 1 AU. A confocal microscopy image of the cells was used to position the crosshairs of the microscope laser in the nucleus of 10-15 cells within the frame. All FCS measurements consisted of ten ten-second traces. A minimum of 30 cells per condition were measured for each biological replicate, and a minimum of two biological replicates were collected for each condition.

The expected diffusion time (*τ*_diff_) for ATTO^TM^ 550-Cas9 RNP was obtained by measuring *in vitro* autocorrelation traces for 400 nM solutions of RNP, annealed ATTO^TM^ 550-tracrRNA:crRNA, or ATTO^TM^ 550-tracrRNA alone in DMEM media (25 mM Hepes, no phenol red) at 37°C (Fig. S6). For each sample, ten ten-second autocorrelation traces using the settings described above were measured per point, and three points were obtained per sample.Given the distribution of τ_diff_ values measured for free RNA in buffer versus in cells, we employed a lower τ_diff_ filter of 0.5 ms for Cas9 RNP delivery experiments to avoid fluorescent signal contributed by free RNA. These data were fitted using eq (1) below to derive the average diffusion time (τ_diff_).

### Analysis of FCS Data

Autocorrelation traces obtained from FCS measurements were analyzed using a custom MATLAB script.(40, 41) To extract quantitative information from *in cellula* data, the effective confocal volume of the microscope must be known. This value was determined using eqs 1-3 by and the *in vitro* autocorrelation traces for the AlexaFluor 594 hydrazide standard measured at the start of each experiment, which have known diffusion coefficients in water. (42, 43).These traces were fitted to a 3D diffusion equation (eq 1):

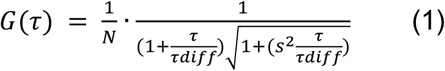

where N = the average number of molecules detected in the focal volume (V_eff_), *τ*_diff_ = the average diffusion time that a molecule requires to cross V_eff_, and s = the structure factor (the ratio of the radial to axial dimensions of the focal volume). The structure factor was measured to be 0.17 using the autocorrelation function of AlexaFluor 594 in water at 25°C and fixed for all subsequent analysis. V_eff_ can be extracted from these data by inserting the *τ*_diff_ value derived from eq (1) to calculate ω_1_ in eq (2):

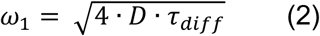

where ω_1_ = the lateral extension of the confocal volume and D = the known diffusion coefficient of the dye standard in water at 37°C. Note that diffusion coefficients at 25°C are typically reported in the literature and can be used to calculate the diffusion coefficient at 37°C using eq (3):

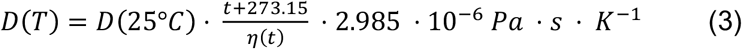

where t = 37°C, D(25°C) for AlexaFluor 594 is 3.88 • 10^-6^cm^2^/s and for AlexaFluor 488 is 4.14 • 10^-6^cm^2^/s (ref.^47^), and q(t) is the viscosity of water at 37°C (6.913 • 10^-4^ Pa • s, ref.^30^). Using this formula, the diffusion coefficient of AlexaFluor 594 at 37° is 5.20 • 10^-6^ cm^2^/s and that of AlexaFluor 488 is 5.54 • 10’^6^ cm^2^/s.

V_eff_ can then be directly calculated from eq (2):

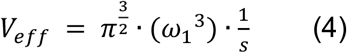

The average V_eff_ for all experiments ranged from 0.25-0.4 fL.

To determine the appropriate fitting equation for data collected in cells, autocorrelation traces derived from *in cellula* measurements of HeLa cells treated with 0.24×10^7^ to 15×10^7^ Cas9 per cell were fitted using two equations. The first was a 3D anomalous diffusion equation used for previous FCS measurements of proteins in live cells (eq 5)(39):

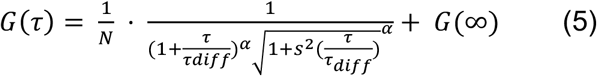

where N = the average number of molecules in the focal volume, *τ*_diff_ = the average diffusion time that a molecule requires to cross V_eff_, α = the anomalous diffusion coefficient, and s = the structure factor (0.17).

The second equation was a two-component diffusion equation previously applied to analysis of DNA-binding transcription factors by FCS.(41, 44). The equation incorporates both a rapidly and slowly diffusing component to account for biphasic autocorrelation functions. This equation is almost identical to a two-component diffusion equation used for previous single-molecule analysis of Cas9 in live cells, (45) except it incorporates an anomalous diffusion coefficient for the slow-diffusing fraction. (41) Since Cas9 binds DNA, we expected eq (6) to more accurately fit ACFs in live cells than eq 5:

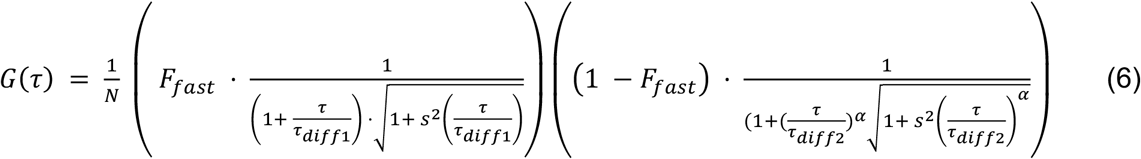

where N = the average number of molecules in the focal volume, *τ*_diff1_ = the average diffusion time for the rapidly-diffusing component, *τ*_diff2_ = the average diffusion time for the slow-diffusing component, α = the anomalous diffusion coefficient, F_fast_ = the fraction of molecules that are rapidly diffusing, and s = the structure factor (0.17).

From the fittings, a set of parameters specific to individual measurements was obtained, including the diffusion time of the detected molecules (a single τ_diff_ for the 1-component fit or *τ*_diff1_ and *τ*_diff2_ for the 2-component fit), the fraction of molecules rapidly diffusing (F_fast_), the number of molecules detected in the focal volume, and a chi-square (χ^2^) value to describe the goodness of fit. The χ^2^ values were compared for the one-component vs two-component fits to determine that the two-component diffusion equation most accurately represents the data. All FCS data were therefore fitted by eq (6) and filtered as described previously (41) to remove measurements where *τ*_diff1_ < 0.5 ms (indicative of free RNA), *τ*_diff1_ > 10 ms (indicative of aggregation), α < 0.3, and χ^2^ > 30. We also excluded fits for which a second component was not identified.

For all FCS curves that passed these thresholds, the concentration (C) of protein in the nucleus was then calculated using the value of N derived above in eq (7):

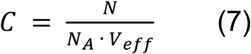

where N_A_ = Avogadro’s number (6.023 × 10^23^ mol^-1^). At least 20 concentration values from curves that passed all filters were used for each FCS condition.

The number of Cas9 molecules in the nucleus was calculated using the concentration obtained from eq (7) and a HeLa nuclear volume of 6.90 • 10^-13^ L (ref.^38^) using eq (8):

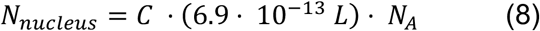

where N_A_ = Avogadro’s number (6.023 × 10^23^ mol^-1^).

Finally, each concentration value derived from eq (7) had a corresponding F_fast_ value describing the fraction of this concentration that was rapidly diffusing. The concentration of DNA-bound Cas9 in the nucleus was then calculated using eq 9:

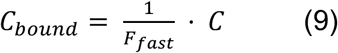

### Cas9 Molecules per Nucleus Calculations

The number of Cas9 molecules per nucleus was calculated by multiplying the average volume of the HeLa nuclei, 690 μM^3^ (46), by the nuclear concentration determined by FCS. To estimate the number of Cas9 RNPs per nucleus that are typically required for productive editing by nucleofection we first estimated the total Cas9 doses required for editing by electroporation by analyzing the Hela dose curves (Fig. S3B). The median Cas9 RNP dosage for half maximal (EC50) editing by electroporation was 6.4×10^7^ Cas9 per cell (Fig. S3B). The high efficiency B2M guide (Supplementary table 3) EC50 was 7.2×10^6^ Cas9 per cell in HeLa cells. Using the linear regression from the FCS electroporation dose titration (Fig. 2D, Supplementary Table 2), we calculated that this dosage would amount to nuclear concentration of 3.2 nM. Using Avogadro’s number (6.023 × 10^23^ mol^-1^), this amounts to ∼1300 Cas9 RNP molecules per nucleus.

### Western Blotting and densitometry

Samples were denatured by mixing with 5x Laemmli with 10% 2-mercaptoethanol and heating at 95°C for 3 minutes. Samples were run on 4%–20% SDS-PAGE gels (Bio-Rad) prior to transfer onto a methanol soaked polyvinylidene difluoride (PVDF, Bio-Rad) membrane. PVDF membranes were blocked with 10% non-fat milk (Apex) in 1xPBS (GIBCO) with 0.1% Tween (Sigma) (PBS-T) for one hour at room temperature (∼22-25°C). The solution was replaced with 0.1% non-fat milk in PBS-T and primary antibody dilution (Supplementary table 1) in 1% non-fat milk in PBS-T incubated at 4°C overnight. The following day, the solution was replaced with 1% non-fat milk in PBS-T and a secondary antibody dilution (Supplementary table 1) and gently shaken for 1 hour. Western blot membranes were washed with PBS-T three times, with 2-3 minutes wash steps, prior to imaging on a LI-COR OdysseyCLx. Fiji (previously imageJ) was used to quantify relative band intensity on western blots.

### Quantification of Cas9 RNPs per EDV

The Cas9 ELISA kit (Cell BioLabs Inc.) and lenti-X p24 Rapid titer kit (Takara Biosciences) were used to quantify the cas9 and p24 in Cas9-EDVs, respectively. For cas9 and p24 measurements, Cas9-EDVs were diluted 20-2000 fold and 1000-100,000 fold, respectively. Both ELISA were done according to the manufacturer’s protocol. Absorbance at 450 nm was measured by a plate reader (Biotek). The amount of Cas9 in the samples was determined by comparison to serial dilution to a cas9 standard (Cell Biolabs Inc.). The amount of p24 was determined by comparison to serial dilution to a p24 standard (Takara Biosciences). The number of p24 per EDV was approximated to be 2500 CA molecules per particle (47).

Cas9 from lyzed EDVs were measured by ELISA and validated against purified Cas9 from commercial (IDT, Cell Biolabs) and in-house sources (UC-Berkeley, Macrolab, QB3), quantified with photospectrometry (Fig. S2).

Quantitative RT-PCR of B2M sgRNA in EDVs

Quantitative RT-PCR was done as previously published(18). Cas9-EDVs containing the sgRNA targeting the B2M gene were produced and concentrated as described above. RNA was extracted from 150 ul of Cas9-EDVs using the NucleoSpin RNA Virus Kit (Takara Bio) following the manufacturer’s instructions. Quantitative RT-PCR was done with PrimeTimeTM One-Step RT-qPCR Master Mix (IDT) following the manufacturers instructions for the QuantStudio 6 Flex Real-Time PCR system (Thermo Fisher). The qPCR primers were custom ordered as a TaqMan Small RNA Assay to detect the B2M sgRNA (Supplementary Table 3)

### In vitro cleavage assessment of Cas9 activity

Prior to use, double stranded DNA (dsDNA) substrate (Supplemental table 1) was annealed in 10 mM HEPES, 20 mM KCl, 1.5 mM MgCl_2_ at 95 °C for 5 minutes and cooled at a rate of 1 C/min to 4C. The annealed substrate was run on 8% Native-PAGE and annealed band was excised, ground finely, and eluted in 5 ml of water overnight at 4C. Next day, this solution was filtered through a 0.22 uM filter, concentrated with 3 kDa spin filter (Amicon), and ethanol precipitated. The dried pellet was resuspended with DEPC water, and concentration was determined by spectrophotometry.

Single guide targeting the spacer was complexed with Cas9 protein in 2:1 ratio for 15 minutes at 37C in 200 mM HEPES; 1M KCl; 100 mM MgCl2, 10% Glycerol, 5 mM DTT. Following complexation, the RNP was diluted and added in 1:1 stoichiometry to annealed dsDNA substrate (Supplemental table 1) at final concentrations of 100 nM. The cleavage reaction occurred at 37 °C for 2 hrs in 20 mM HEPES; 100 mM KCl; 10 mM MgCl2, 1% Glycerol, 0.5 mM DTT.

### Next Generation Sequencing (NGS) to assess genome editing

Next-generation sequencing was used for detection of on-target genome editing in 293T cells. Genomic DNA was extracted using QuickExtract (Lucigen) as previously described (17). Q5 high fidelity polymerase (NEB) was used to attach adapters to the Cas9-RNP target site amplicons (Supplemental table 1). The resulting PCR1 products were cleaned up using magnetic SPRI beds (UC Berkeley DNA Sequencing Facility). Library preparation and sequencing was performed by the Innovative Genomics Institute Next Generation Sequencing Core using MiSeq v2 (Illumina). Reads were analyzed with CRISPResso2 (http://crispresso.pinellolab.partners.org/login).

### Digital Droplet quantitative PCR (ddPCR)

Cells were collected at day 4, unless otherwise stated in figures, following editing with electroporation or EDVs, and genomic DNA was extracted with QuickExtract DNA Extraction Solution (Lucigen).

For DSB detection, the ddPCR setup was similar to what has been previously described (48, 49), with two ∼200 bp amplicons for the *B2M* target gene (Supplementary Table 3). Amplicon 1 was located proximal to the centromere and utilized a 5’ hexachlorofluorescein-labeled (HEX) oligonucleotide probe (PrimeTime qPCR probes, Zen double quencher, IDT). Amplicon 2 was located ∼200 bp away from amplicon 1, and utilized a 5’ 6-fluorescein-labeled (FAM) oligonucleotide probe (PrimeTime qPCR probes, Zen double quencher, IDT). Amplicon 1 served as a reference that should be unaffected by Cas9 genome editing and would signal whether *B2M* was in a given droplet. Amplicon 2 spanned the *B2M* target site, with the probe located ∼50 bp from the cleavage site. If the target site was not repaired after Cas9 cleavage, or if the chromosome was lost(48), amplicon 2 would not be amplified and the FAM probe would be quenched. ddPCR reactions were assembled with ddPCR Supermix for Probes (No dUTP, Bio-Rad), 900 nM of each primer, 250 nM of each probe, and 15-30 ng gDNA.

Droplets were formed using a QX200 Droplet Generator (Bio-Rad) following the manufacturer’s instructions prior to PCR. The next day, ddPCR droplets were analyzed on a QX200 Droplet Reader (Bio-Rad). Data was analyzed with the QX Manager Software (Bio-Rad), and thresholds were set manually based on wells with untreated reference samples. The percentage of DSBs was calculated based on droplets that had the reference amplicon 1 (HEX+) but did not produce the neighboring amplicon (FAM+) (equation below).

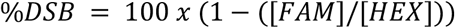

### Immunofluorescent (IF) Imaging and Quantification

For imaging EDV delivery in HeLa cells, 400k cells were plated the evening prior in a 12 cm dish that had 18 mM coverslips precoated with poly-L-lysine (Sigma-Aldrich #P7886, 100 ug/ml in 75 mM NaCl, 50 mM Na_2_H_20_B_4_O_17_, pH 8.4) For EDV experiments, unless otherwise stated in figure legends, 350 ul of 20x concentrated EDV mixture were added to the cells in 1:1 mix with OptiMEM (final volume 700 ul). EDV containing media was swapped for prewarmed supplemented DMEM (see above) at ∼4 hours. For imaging Cas9 delivered by electroporation, 3 million cells were electroporated with 600 pmol RNP (1.2×10^8^ Cas9 per cell). Following delivery, at timepoints specified in the figures, cells were fixed with 4% PFA in 1xDPBS for 10 min (ThermoScientific #28908). For time course experiments, all cells were fixed and then stored in 1xDPBS until all samples were harvested, so that permeabilization and staining steps were done simultaneously. We then washed coverslips three times with 1×DPBS, and permeabilized samples with 0.2% Triton-X-100 in 1xDPBS for 10 minutes. Following permeabilization, samples were washed three times, and then blocked with Image-iT FX signal enhancer (Invitrogen). After this initial blocking step, samples were washed twice with 1xDPBS and then further blocked in 10% Goat Serum (ThermoFisher #50062Z) for 20-30 minutes. Then samples were incubated with primary antibodies (Supplementary table 1) diluted in 10% Goat Serum at 4C overnight. The following day, the coverslips were washed three times with 1xDPBS and incubated for 1 hour with secondary antibodies (Supplementary Table 1). For EDV confocal microscopy, secondary antibodies from tyramide signal superboost kits (ThermoFisher, Supplementary Table 1) were used following manufacturers instructions. The labeling reaction was done for 2 minutes for all EDV experiments. Samples were then washed three times and mounted with Prolong Diamond antifade mountant (Invitrogen). All slides were stored at -20 C for periods longer than 1 week. All fixed cell confocal microscopy was performed using the Zeiss LSM710 microscope (UC-Berkeley, Bioimaging Facility).

For quantification of relative nuclear intensity of Cas9, all Z-stacks were obtained based off of half the wavelength of emission of the Cas9 associated fluorophore (244 nm, Nyquist sampling) through the Z-plane of the HeLa cells that were counterstained with SYTOX™ Deep Red Nucleic Acid Stain (1:2000, Invitrogen, #S11380). These images were deconvoluted using the default settings of Huygens Professional (v22.10) to reduce noise. Using Imaris (v10.2) the entire nuclear counterstained region was outlined and the median nuclear intensity in the 488 channel (Cas9 channel) was quantified. Any nuclei that were on the edges of the image, were actively dividing, or that could not be independently quantified due to their close proximity to other nuclei, were manually excluded from quantification.

### Flow Cytometry

Cells were stained with anti-human B2M-PE (316306, Biolegend) in PBS containing 1% bovine serum albumin. An Attune NxT flow cytometer equipped with a 96-well autosampler (Thermo Fisher Scientific) was used for flow cytometry acquisition. Data analysis was performed using FlowJo v10 10.7.1 (FlowJo, LLC, Ashland OR).

### Statistical analysis

Statistical analysis was performed using Prism v10 unless otherwise stated. Statistical details for experiments, including the values and definitions of the samples sizes and error bars are reported in the figure legends. Unless otherwise specified in figure legend, two sided t-tests were used for pairwise comparisons and ANOVA was used for multiple comparisons.

## Data and materials availability

Flow cytometry, fluorescence correlation spectroscopy, unprocessed ddPCR data, quantified confocal Z-stacks and next generation sequencing (NGS) raw files are available upon request. All other data is in the main text or supplementary materials.

## Acknowledgements

We appreciated useful discussions with Ross Wilson, Laura Hofman, Matthew Kan and Kai Chen. We thank Matthew Kan for assisting in setting up the ddPCR assay. H.K. was in part supported by the National Institutes of Health (NIH) Stem cell biological engineering training program (Grant no. T32GM098218). M.Z. was in part supported by the National Science Foundation (Grant no. 2203903) and the NIH (Grant no. R35GM134963). W.N .was supported by Natural Sciences and Engineering Research Council of Canada Postdoctoral Fellowship PDF-578176-2023. Microscopy research reported in this publication was supported in part by the NIH S10 program under the award number 1S10RR026866-01. The content is solely the responsibility of the authors and does not necessarily represent the official views of the National Institutes of Health.

## References

1. Raguram, A., Banskota, S. and Liu, D.R. (2022) Therapeutic in vivo delivery of gene editing agents. Cell, 185, 2806–2827.

2. Sinclair, F., Begum, A.A., Dai, C.C., Toth, I. and Moyle, P.M. (2023) Recent advances in the delivery and applications of nonviral CRISPR/Cas9 gene editing. Drug Deliv. Transl. Res., 13, 1500–1519.

3. Jang, H.-K., Jo, D.H., Lee, S.-N., Cho, C.S., Jeong, Y.K., Jung, Y., Yu, J., Kim, J.H., Woo, J.-S. and Bae, S. (2021) High-purity production and precise editing of DNA base editing ribonucleoproteins. Sci Adv, 7.

4. Wienert, B., Shin, J., Zelin, E., Pestal, K. and Corn, J.E. (2018) In vitro-transcribed guide RNAs trigger an innate immune response via the RIG-I pathway. PLoS Biol., 16, e2005840.

5. Kim, S., Koo, T., Jee, H.-G., Cho, H.-Y., Lee, G., Lim, D.-G., Shin, H.S. and Kim, J.-S. (2018) CRISPR RNAs trigger innate immune responses in human cells. Genome Res., 28, 367–373.

6. Charlesworth, C.T., Deshpande, P.S., Dever, D.P., Camarena, J., Lemgart, V.T., Cromer, M.K., Vakulskas, C.A., Collingwood, M.A., Zhang, L., Bode, N.M., et al. (2019) Identification of preexisting adaptive immunity to Cas9 proteins in humans. Nat. Med., 25, 249–254.

7. van Haasteren, J., Li, J., Scheideler, O.J., Murthy, N. and Schaffer, D.V. (2020) The delivery challenge: fulfilling the promise of therapeutic genome editing. Nat. Biotechnol., 38, 845–855.

8. Hou, X., Zaks, T., Langer, R. and Dong, Y. (2021) Lipid nanoparticles for mRNA delivery. Nat. Rev. Mater., 6, 1078–1094.

9. Hendel, A., Bak, R.O., Clark, J.T., Kennedy, A.B., Ryan, D.E., Roy, S., Steinfeld, I., Lunstad, B.D., Kaiser, R.J., Wilkens, A.B., et al. (2015) Chemically modified guide RNAs enhance CRISPR-Cas genome editing in human primary cells. Nat. Biotechnol., 33, 985–989.

10. Hanlon, K.S., Kleinstiver, B.P., Garcia, S.P., Zaborowski, M.P., Volak, A., Spirig, S.E., Muller, A., Sousa, A.A., Tsai, S.Q., Bengtsson, N.E., et al. (2019) High levels of AAV vector integration into CRISPR-induced DNA breaks. Nat. Commun., 10, 4439.

11. Espinoza, D.A., Fan, X., Yang, D., Cordes, S.F., Truitt, L.L., Calvo, K.R., Yabe, I.M., Demirci, S., Hope, K.J., Hong, S.G., et al. (2019) Aberrant Clonal Hematopoiesis following Lentiviral Vector Transduction of HSPCs in a Rhesus Macaque. Mol. Ther., 27, 1074–1086.

12. Porello, I. and Cellesi, F. (2023) Intracellular delivery of therapeutic proteins. New advancements and future directions. Front. Bioeng. Biotechnol., 11, 1211798.

13. Frangoul, H., Altshuler, D., Cappellini, M.D., Chen, Y.-S., Domm, J., Eustace, B.K., Foell, J., de la Fuente, J., Grupp, S., Handgretinger, R., et al. (2021) CRISPR-Cas9 gene editing for sickle cell disease and β-thalassemia. N. Engl. J. Med., 384, 252–260.

14. Hanna, R., Frangoul, H., Mckinney, C., Pineiro, L., Mapara, M., Chang, K.-H., Jaskolka, M., Kim, K., Rizk, M., Afonja, O., et al. (2023) S264: Edit-301 shows promising preliminary safety and efficacy results in the phase i/ii clinical trial (Ruby) of patients with severe sickle cell disease using highly specific and efficient ascas12a enzyme. HemaSphere, 7, e05170e0.

15. Kim, S., Kim, D., Cho, S.W., Kim, J. and Kim, J.-S. (2014) Highly efficient RNA-guided genome editing in human cells via delivery of purified Cas9 ribonucleoproteins. Genome Res., 24, 1012–1019.

16. Foss, D.V., Muldoon, J.J., Nguyen, D.N., Carr, D., Sahu, S.U., Hunsinger, J.M., Wyman, S.K., Krishnappa, N., Mendonsa, R., Schanzer, E.V., et al. (2023) Peptide-mediated delivery of CRISPR enzymes for the efficient editing of primary human lymphocytes. Nat Biomed Eng, 7, 647–660.

17. Hamilton, J.R., Tsuchida, C.A., Nguyen, D.N., Shy, B.R., McGarrigle, E.R., Sandoval Espinoza, C.R., Carr, D., Blaeschke, F., Marson, A. and Doudna, J.A. (2021) Targeted delivery of CRISPR-Cas9 and transgenes enables complex immune cell engineering. Cell Rep., 35, 109207.

18. Hamilton, J.R., Chen, E., Perez, B.S., Sandoval Espinoza, C.R., Kang, M.H., Trinidad, M., Ngo, W. and Doudna, J.A. (2024) In vivo human T cell engineering with enveloped delivery vehicles. Nat. Biotechnol., 10.1038/s41587-023-02085-z.

19. Johnson, N.M., Alvarado, A.F., Moffatt, T.N., Edavettal, J.M., Swaminathan, T.A. and Braun, S.E. (2021) HIV-based lentiviral vectors: origin and sequence differences. Mol. Ther. Methods Clin. Dev., 21, 451–465.

20. Strebinger, D., Frangieh, C.J., Friedrich, M.J., Faure, G., Macrae, R.K. and Zhang, F. (2023) Cell type-specific delivery by modular envelope design. Nat. Commun., 14, 5141.

21. Leibowitz, M.L., Papathanasiou, S., Doerfler, P.A., Blaine, L.J., Sun, L., Yao, Y., Zhang, C.-Z., Weiss, M.J. and Pellman, D. (2021) Chromothripsis as an on-target consequence of CRISPR-Cas9 genome editing. Nat. Genet., 53, 895–905.

22. Kim, S.A., Heinze, K.G. and Schwille, P. (2007) Fluorescence correlation spectroscopy in living cells. Nat. Methods, 4, 963–973.

23. Elson, E.L. (2011) Fluorescence correlation spectroscopy: past, present, future. Biophys. J., 101, 2855– 2870.

24. Schwille, P. (2001) Fluorescence correlation spectroscopy and its potential for intracellular applications. Cell Biochem. Biophys., 34, 383–408.

25. Moon, S.B., Kim, D.Y., Ko, J.-H., Kim, J.-S. and Kim, Y.-S. (2019) Improving CRISPR genome editing by engineering guide RNAs. Trends Biotechnol., 37, 870–881.

26. Lazzarotto, C.R., Malinin, N.L., Li, Y., Zhang, R., Yang, Y., Lee, G., Cowley, E., He, Y., Lan, X., Jividen, K., et al. (2020) CHANGE-seq reveals genetic and epigenetic effects on CRISPR-Cas9 genome-wide activity. Nat. Biotechnol., 38, 1317–1327.

27. Wu, X., Scott, D.A., Kriz, A.J., Chiu, A.C., Hsu, P.D., Dadon, D.B., Cheng, A.W., Trevino, A.E., Konermann, S., Chen, S., et al. (2014) Genome-wide binding of the CRISPR endonuclease Cas9 in mammalian cells. Nat. Biotechnol., 32, 670–676.

28. Doench, J.G., Fusi, N., Sullender, M., Hegde, M., Vaimberg, E.W., Donovan, K.F., Smith, I., Tothova, Z., Wilen, C., Orchard, R., et al. (2016) Optimized sgRNA design to maximize activity and minimize off-target effects of CRISPR-Cas9. Nat. Biotechnol., 34, 184–191.

29. Moreno-Mateos, M.A., Vejnar, C.E., Beaudoin, J.-D., Fernandez, J.P., Mis, E.K., Khokha, M.K. and Giraldez, A.J. (2015) CRISPRscan: designing highly efficient sgRNAs for CRISPR-Cas9 targeting in vivo. Nat. Methods, 12, 982–988.

30. Liu, M.-S., Gong, S., Yu, H.-H., Jung, K., Johnson, K.A. and Taylor, D.W. (2020) Engineered CRISPR/Cas9 enzymes improve discrimination by slowing DNA cleavage to allow release of off-target DNA. Nat. Commun., 11, 3576.

31. Gong, S., Yu, H.H., Johnson, K.A. and Taylor, D.W. (2018) DNA Unwinding Is the Primary Determinant of CRISPR-Cas9 Activity. Cell Rep., 22, 359–371.

32. Shi, H., Al-Sayyad, N., Wasko, K.M., Trinidad, M.I., Doherty, E.E., Vohra, K., Boger, R.S., Colognori, D., Cofsky, J.C., Skopintsev, P., et al. (2024) Rapid two-step target capture ensures efficient CRISPR-Cas9-guided genome editing. bioRxiv, 10.1101/2024.10.01.616117.

33. Chen, K., Han, H., Zhao, S., Xu, B., Yin, B., Trinidad, M., Burgstone, B.W., Murthy, N. and Doudna, J.A. (2023) Lung and liver editing by lipid nanoparticle delivery of a stable CRISPR-Cas9 RNP. bioRxivorg, 10.1101/2023.11.15.566339.

34. Kotnik, T., Rems, L., Tarek, M. and Miklavčič, D. (2019) Membrane electroporation and electropermeabilization: Mechanisms and models. Annu. Rev. Biophys., 48, 63–91.

35. Ci, Y., Yang, Y., Xu, C. and Shi, L. (2018) Vesicular stomatitis virus G protein transmembrane region is crucial for the hemi-fusion to full fusion transition. Sci. Rep., 8, 10669.

36. Chen, J., Su, S., Pickar-Oliver, A., Chiarella, A.M., Hahn, Q., Goldfarb, D., Cloer, E.W., Small, G.W., Sivashankar, S., Ramsden, D.A., et al. (2024) Engineered Cas9 variants bypass Keap1-mediated degradation in human cells and enhance epigenome editing efficiency. Nucleic Acids Res.

37. Ramadoss, G.N., Namaganda, S.J., Hamilton, J.R., Sharma, R., Chow, K.G., Macklin, B.L., Sun, M., Liu, J.-C., Fellmann, C., Watry, H.L., et al. (2024) Neuronal DNA repair reveals strategies to influence CRISPR editing outcomes. bioRxivorg, 10.1101/2024.06.25.600517.

38. Ngo, W., Peukes, J.T., Baldwin, A., Xue, Z.W., Hwang, S., Stickels, R.R., Lin, Z., Satpathy, A.T., Wells, J.A., Schekman, R., et al. (2024) Mechanism-guided engineering of a minimal biological particle for genome editing. bioRxivorg, 10.1101/2024.07.23.604809.

39. Knox, S.L., Steinauer, A., Alpha-Cobb, G., Trexler, A., Rhoades, E. and Schepartz, A. (2020) Chapter Twenty-One - Quantification of protein delivery in live cells using fluorescence correlation spectroscopy. In Chenoweth, D.M. (ed), Methods in Enzymology. Academic Press, Vol. 641, pp. 477–505.

40. Zoltek, M., Vázquez Maldonado, A.L., Zhang, X., Dadina, N., Lesiak, L. and Schepartz, A. (2024) HOPS-Dependent Endosomal Escape Demands Protein Unfolding. ACS Cent Sci, 10, 860–870.

41. Zhang, X., Cattoglio, C., Zoltek, M., Vetralla, C., Mozumdar, D. and Schepartz, A. (2023) Dose-Dependent Nuclear Delivery and Transcriptional Repression with a Cell-Penetrant MeCP2. ACS Cent Sci, 9, 277– 288.

42. Steinauer, A., LaRochelle, J.R., Knox, S.L., Wissner, R.F., Berry, S. and Schepartz, A. (2019) HOPS-dependent endosomal fusion required for efficient cytosolic delivery of therapeutic peptides and small proteins. Proc. Natl. Acad. Sci. U. S. A., 116, 512–521.

43. Petrov, E.P. and Schwille, P. (2008) State of the art and novel trends in fluorescence correlation spectroscopy. In Springer Series on Fluorescence. Springer Berlin Heidelberg, Berlin, Heidelberg, pp. 145–197.

44. Kaur, G., Costa, M.W., Nefzger, C.M., Silva, J., Fierro-González, J.C., Polo, J.M., Bell, T.D.M. and Plachta, N. (2013) Probing transcription factor diffusion dynamics in the living mammalian embryo with photoactivatable fluorescence correlation spectroscopy. Nat. Commun., 4, 1637.

45. Knight, S.C., Xie, L., Deng, W., Guglielmi, B., Witkowsky, L.B., Bosanac, L., Zhang, E.T., El Beheiry, M., Masson, J.-B., Dahan, M., et al. (2015) Dynamics of CRISPR-Cas9 genome interrogation in living cells. Science, 350, 823–826.

46. Monier, K., Armas, J.C., Etteldorf, S., Ghazal, P. and Sullivan, K.F. (2000) Annexation of the interchromosomal space during viral infection. Nat. Cell Biol., 2, 661–665.

47. Cimarelli, A. and Darlix, J.-L. (2002) Biomedicine and Diseases: Review¶Assembling the human immunodeficiency virus type 1. Cell. Mol. Life Sci., 59, 1166–1184.

48. Tsuchida, C.A., Brandes, N., Bueno, R., Trinidad, M., Mazumder, T., Yu, B., Hwang, B., Chang, C., Liu, J., Sun, Y., et al. (2023) Mitigation of chromosome loss in clinical CRISPR-Cas9-engineered T cells. Cell, 186, 4567–4582.e20.

49. Rose, J.C., Stephany, J.J., Valente, W.J., Trevillian, B.M., Dang, H.V., Bielas, J.H., Maly, D.J. and Fowler, D.M. (2017) Rapidly inducible Cas9 and DSB-ddPCR to probe editing kinetics. Nat. Methods, 14, 891–896.

50. Sternberg, S.H., Redding, S., Jinek, M., Greene, E.C. and Doudna, J.A. (2014) DNA interrogation by the CRISPR RNA-guided endonuclease Cas9. Nature, 507, 62–67.

