## Supplemental Material for "Packaged delivery of CRISPR-Cas9 ribonucleoproteins accelerates genome editing"

### Supplementary Material

#### Supplemental Figures

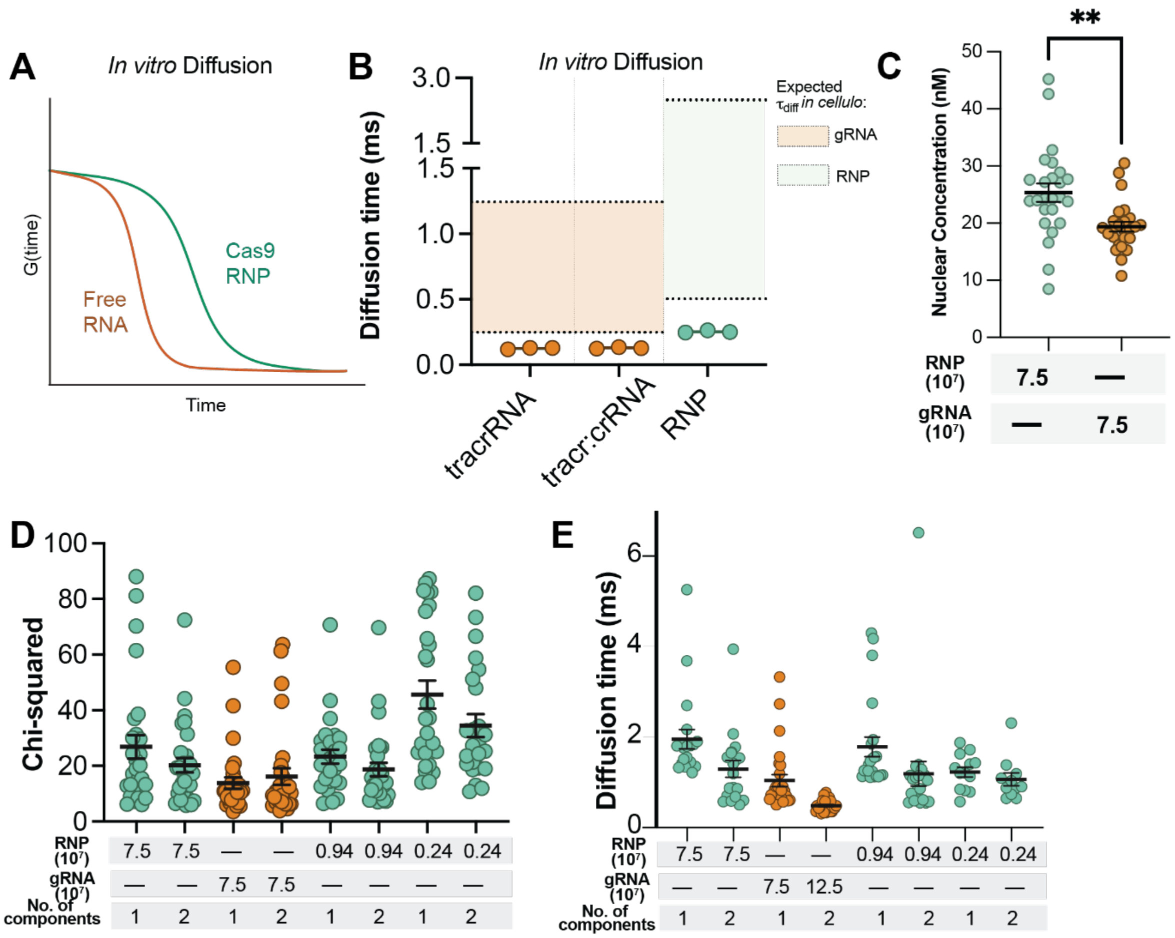


***Supplemental Figure 1: Optimization of fitting parameters for measuring Cas9 RNP nuclear concentration by fluorescence correlation spectroscopy.***

FCS measurements produce an autocorrelation function, which can be fit to derive both the concentration of delivered material and its diffusion time(s). **(A)** Example FCS traces for free sgRNA versus an intact Cas9 RNP. The diffusion time is calculated as the inflection point of the autocorrelation function. **(B)** FCS traces were measured for tracrRNA, tracr:crRNA, and Cas9 RNP *in vitro* (right). Samples were prepared at a final concentration of 400 nM in DMEM. Each point represents the average diffusion time ($t_{diff}$) calculated from ten autocorrelation traces at that point. Average $t_{diff}$ for tracrRNA = 0.126 ms, for sgRNA = 0.129 ms, and for Cas9 RNP = 0.253 ms. **(C)** HeLa cells were electroporated with fluorophore labeled gRNA (ATTO™ 550-tracrRNA:crB2M (IDT)) or with RNP pre-complexed 1:1 with fluorophore labeled guide (7.5 x$10^7$ cas9 per cell). FCS diffusion times provided in ms, n>25 for each FCS condition with at least two biological replicates each (mean ± SEM). **(D)** and **(E)** Comparison of one-component and two-component diffusion fitting equations for FCS data in cells. **(D)** Chi-square ($c^2$) values for autocorrelation curves fitted with one- or two-component diffusion equations plotted as a function of RNP or sgRNA dosage. Each point represents the $c^2$ for the fitting of an average autocorrelation curve (calculated as the mean autocorrelation function from ten individual traces) obtained from an individual cell. **(E)** Diffusion times for the autocorrelation curves fitted in **(D)**. For a one-component diffusion equation, only one diffusion time is produced. For a two-component diffusion equation, a fast diffusion and a slow diffusion time are both calculated; the fast diffusion time is plotted here. Each point corresponds to the $t_{diff}$ derived from each fitted autocorrelation curve in **(D)**.

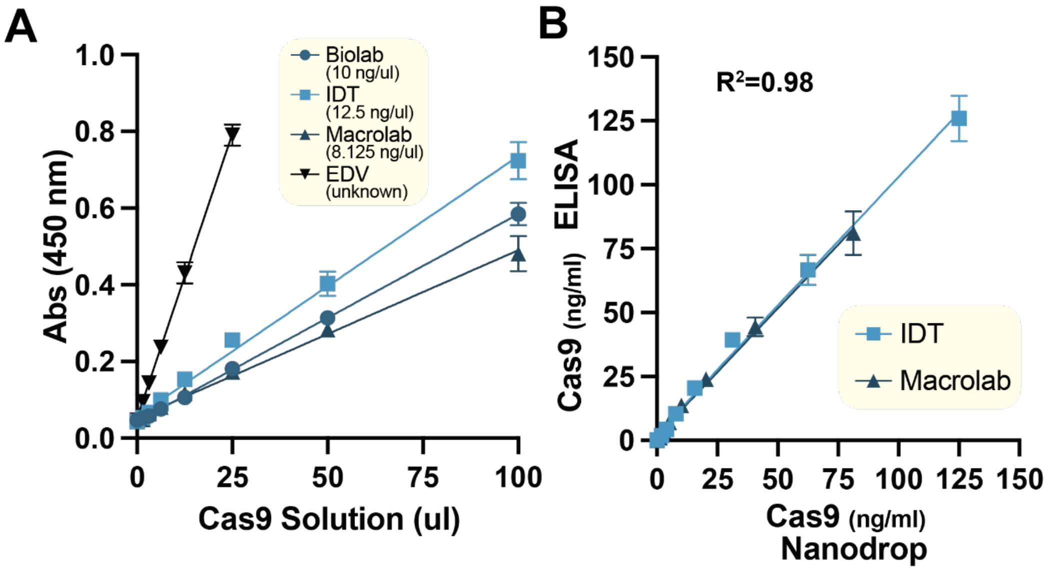


***Supplemental Figure 2: Validation of ELISA for measuring Cas9 used for electroporation and in EDVs***

A) Absorbance values from ELISA for Cas9 as purified RNP diluted 800-fold, from commercial (IDT) and in-house source (UC-Berkeley, Macrolab, QB3), and 20x concentrated EDVs prepared by ultracentrifugation diluted 4-fold (n=3). B) Absorbance values from ELISA used to quantify concentration of Cas9 in commercial (IDT) and in-house (UC-Berkeley, Macrolab, QB3) samples compared to reported concentrations calculated from photospectrometry (Nanodrop) ($R^2$=0.98, n=3).

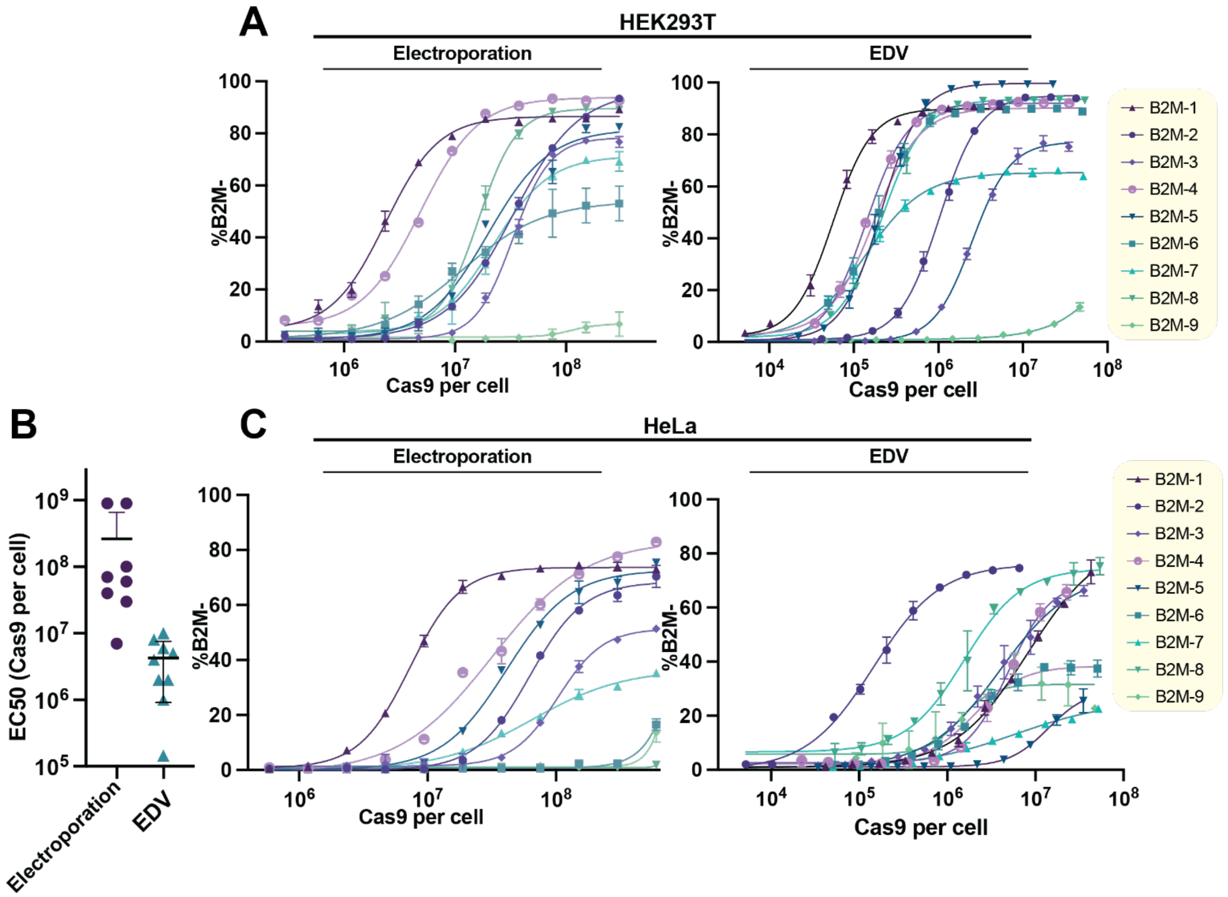


***Supplemental Figure 3: Cas9 RNP doses required for editing by Electroporation and EDVs***

**(A)** Comparison of required RNP doses delivered by electroporation and EDV for 9 different B2M guides in HeLa. Analysis was performed by flow cytometry 4 days post treatment to assess B2M KO. **(B)** HeLa cells treated with Cas9 RNP complexed with the same 9 guides delivered by (Left) electroporation and (Right) EDVs. Cells were analyzed by flow cytometry on day 4 (n=3). **(C)** HEK293T cells were treated with Cas9 RNP complexed with 9 different guides (Supplemental table 1) targeting B2M delivered by (Left) Electroporation and (Right) EDVs. Cells were analyzed by flow cytometry on day 4 (n=3). Datapoints represent the mean with error bars displaying SD. RNP dose curves were modeled (Prism v10) as sigmoidal (4PL, X is concentration).

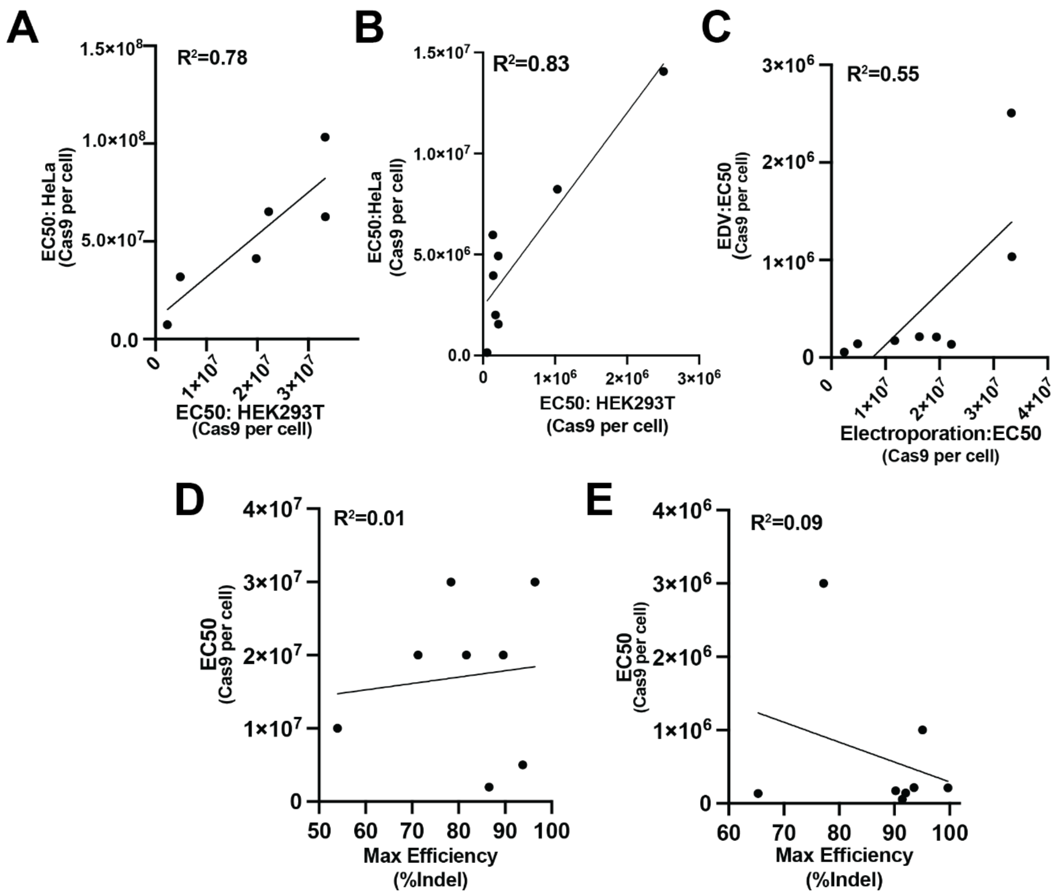


***Supplemental Figure 4: Analyzing the linear relationships between RNP delivery modalities and guide RNA on editing dosage requirements***

Correlation analysis of dosage required for half maximal editing (EC50) from Cas9 RNP delivered by **(A)** Electroporation and **(B)** EDVs in HEK293T (x-axis) and HeLa cells (y-axis).**(C)** Correlation analysis of dosage required for half maximal editing (EC50) from Cas9 RNP delivered by electroporation (x-axis) versus EDV (y-axis). Relationship between gRNA maximum editing efficiency (i.e. the top plateau of dosage curve) versus the dosage EC50 for **(D)** electroporation and **(E)** EDV in HEK293T cells. For correlation analyses, each value calculated from HEK293T HeLa dose curves (Supplemental Figure 3). n=3 technical replicates were used in all experiments.

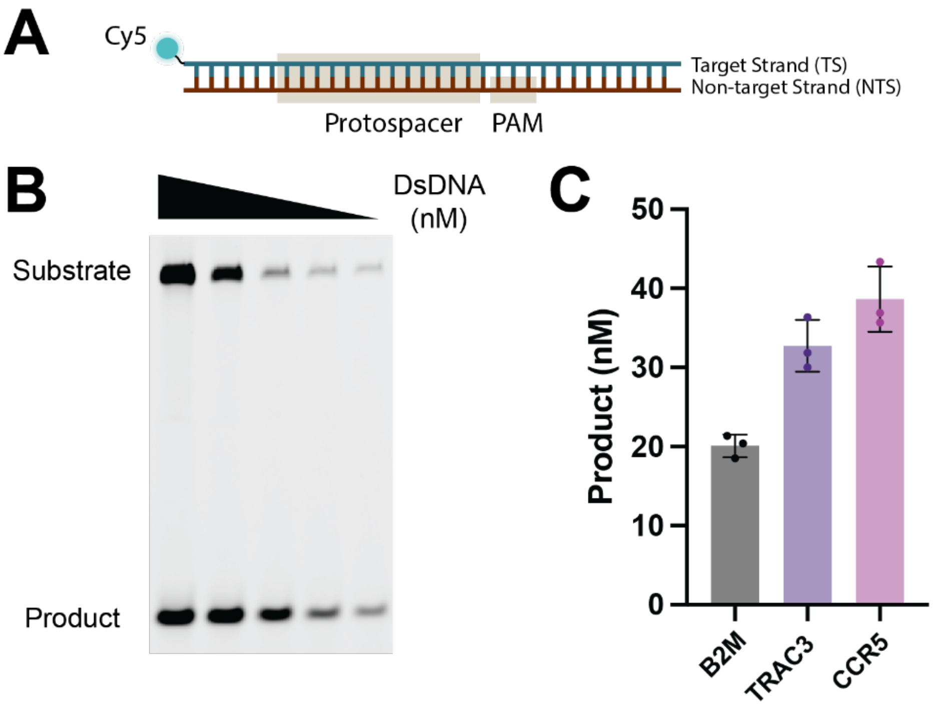


***Supplemental Figure 5: In vitro cleavage assay to assess percent activity of purified Cas9***

**(A)** Schematic of double-stranded DNA (DsDNA) substrate used for assessing the biochemical cleavage activity of purified Cas9 RNP, a single turnover enzyme (50), by active site titration. The Cy5 fluorophore label is on the 5' end of the target strand. **(B)** Representative gel showing 100 nM Cas9 RNP, complexed with TRAC3 guide, incubated with annealed DsDNA substrate with TRAC3 active site at titrated concentrations. Percent cleavage calculated by same method as (31) **(C)** Comparison of cleavage activity of in-house (UC-Berkeley, Macrolab, QB3) Cas9, complexed with respective sgRNA (IDT), targeting substrates with spacers from B2M, TRAC3, CCR5. n=3 technical replicates were used in all experiments. Datapoints represent the mean with error bars displaying SD. Oligos used are in Supplemental table 1.

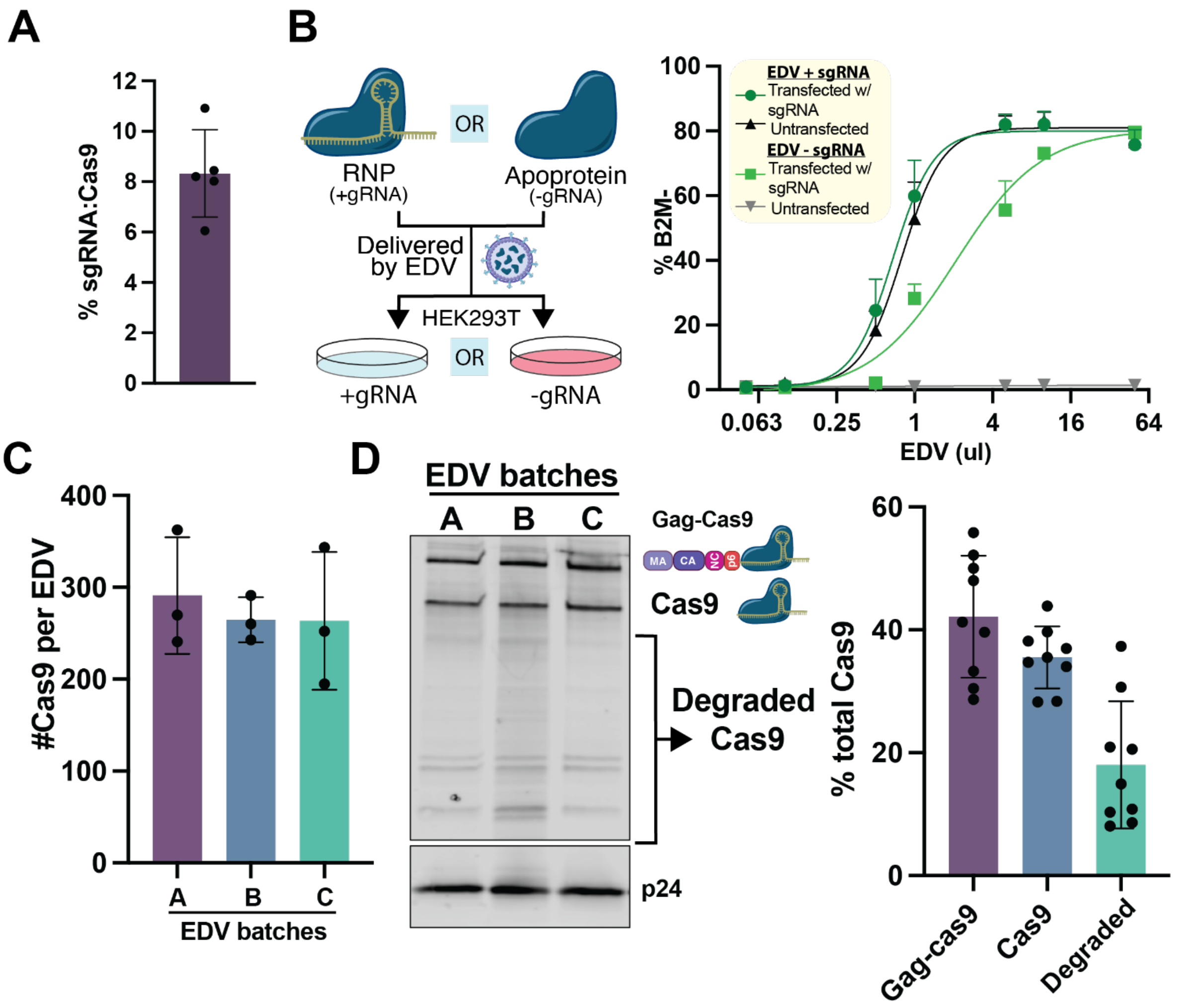


---

***Supplemental Figure 6: Determining the percent of RNPs in EDVs that are functional***

**(A)** The percent of B2M sgRNA available relative to Cas9 protein molecules in EDVs (n=5 biological replicates, mean depicted from 3 technical replicates). The molecules of intact B2M sgRNA and Cas9 protein were determined by quantitative RT-PCR and Cas9 ELISA (n=5 biological replicates measured in triplicate), respectively. **(B)** Schematic of experimental setup to determine whether sgRNA is limiting editing potency of EDVs in cells. HEK293T cells, either untreated or transfected with U6-B2M-sgRNA expression plasmid, were treated with EDVs containing *B2M*-targeting Cas9 RNP or Cas9 apoprotein (n=3). **(C)** B2M knockout was assessed by flow cytometry four days post treatment with minimal difference between conditions where HEK293T cells expressed excess B2M-targeting gRNA versus delivery with

EDVs containing *B2M*-targeting Cas9 RNP. **(D)** Quantification of the number of Cas9 molecules per EDV. p24, the capsid protein and a key structural component of EDVs and native HIV, and Cas9 were quantified by ELISA (n=3 biological replicates measured in triplicate). **(E)** Western blot of 20x ultracentrifuged concentrated EDVs for Cas9 and p24 (loading control). **(F)** Densitometry analysis of the percent of total cas9 in EDVs that remains uncleaved from gag-Cas9, free Cas9, and major Cas9 degradation products.

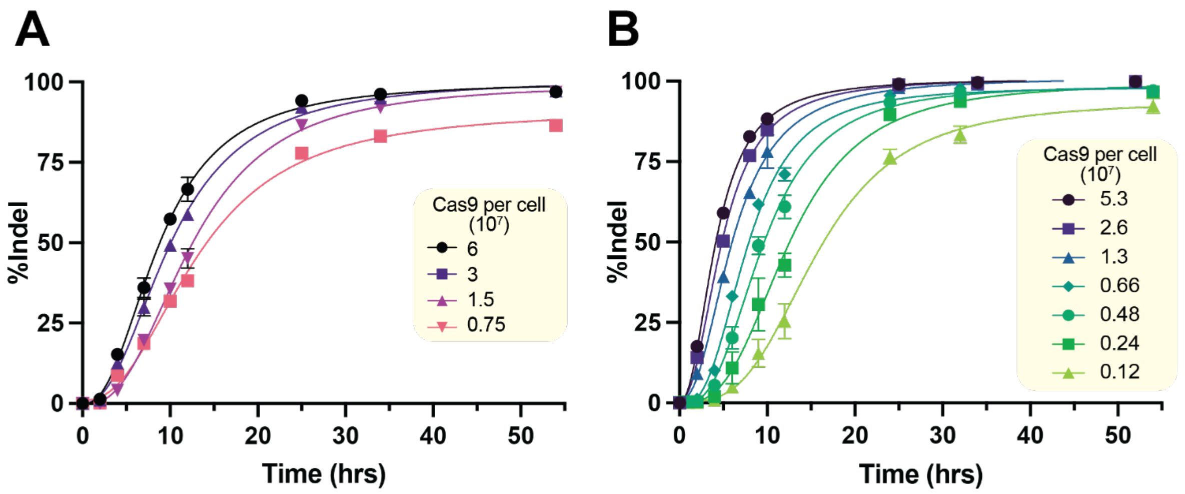


***Supplemental Figure 7: Indels resulting from Cas9 delivery by electroporation and EDVs***

(Relating to Figure 3) Indel formation caused by **(A)** electroporation and **(B)** EDV delivery of *B2M*-targeting RNP in HEK293T cells measured by NGS. All doses are at levels that meet or exceed the amount necessary for 90% maximal editing. The first derivative (rate of indel formation) of these curves is shown in Fig. 3G-H for electroporation and EDV delivery. n=3 technical replicates were used in all experiments. Datapoints represent the mean with error bars displaying SD.

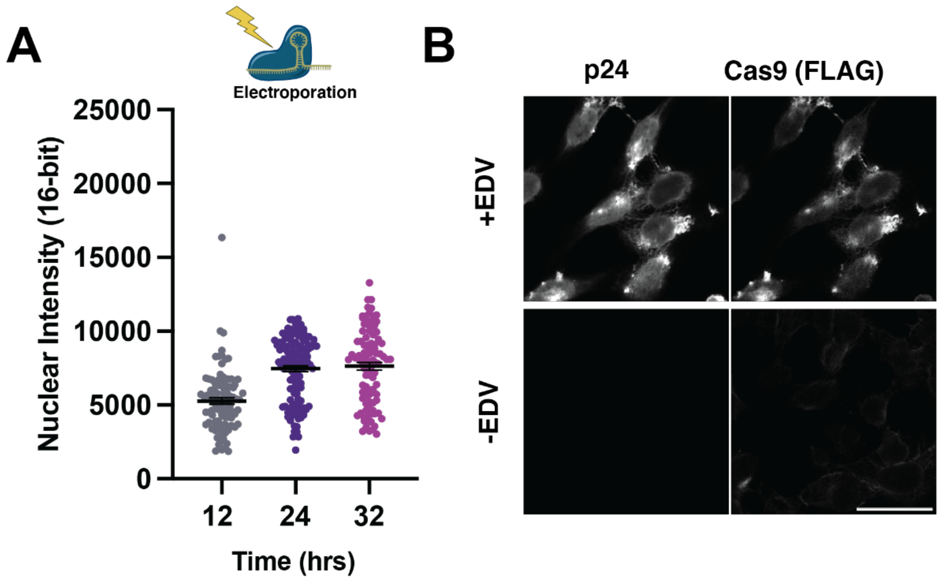


***Supplemental Figure 8: Fixed cell confocal microscopy for relative quantification of Cas9 RNP***

**(A)** Relative quantification of median nuclear intensity of Cas9 delivered by electroporation at 12, 24 and 36 hours. (Representative images in Fig. 4B). Intensity on 16-bit scale (0 to 65536). Each data point represents an individual nuclei n> 94 (mean ± SEM). (see Methods for details).

**(B)** Validation of detection specificity for Cas9 and p24, a HIV capsid protein found in EDVs, stained in cells incubated with EDVs (350 ul 20x ultracentrifuged EDV, 1:1 dilution in OptiMEM) or OptiMEM media for 4 hours, then grown in supplemented DMEM until harvest at 32 hours (see Methods for details). Both EDV treated and untreated cells were stained with both primary and secondary antibodies (Supplemental Table 3).

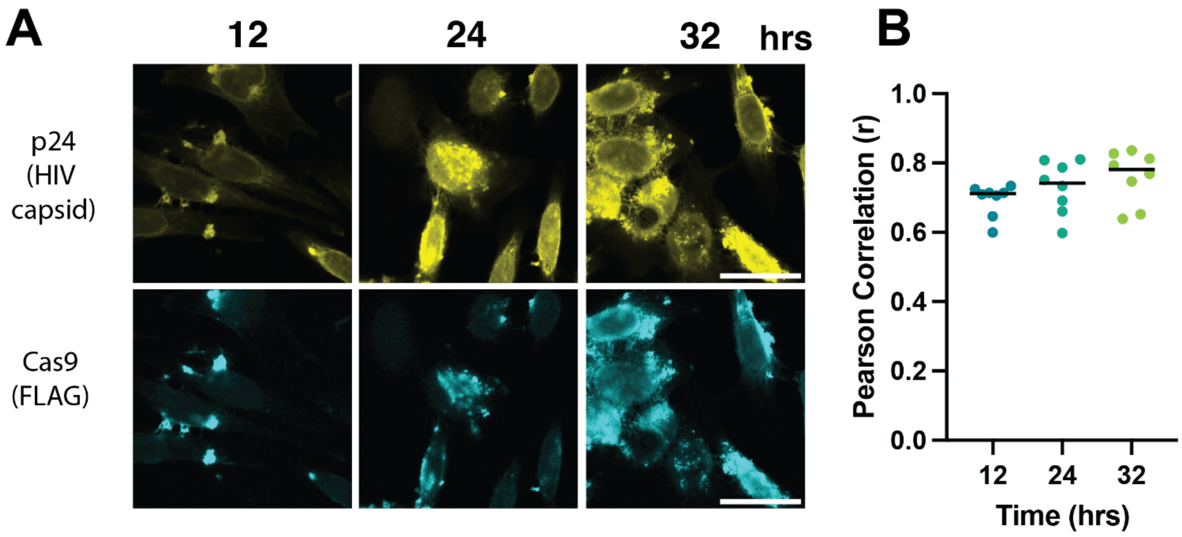


***Supplemental Figure 9: Cas9 in EDV remains uncleaved from Gag, HIV structural polyprotein, in vitro and in cells***

**(C)** Representative images of EDV treated cells at 12, 24 and 36 hours stained to image p24 (Above) and Cas9 (Below), showing consistent colocalization of Cas9 and Gag. **(D)** Quantification of the colocalization of p24, a structural part of gag used in EDV packaging, and Cas9 at 12, 24 and 36 hours. Each data point represents an independent z-stack (n=8) quantified with Pearson's correlation analysis (HuygensPro, see Methods for details).

***Supplementary Table 2: Summary Values for in cellula FCS Experiments***

For all fluorescence correlation spectroscopy (FCS) measurements, *n-BR* refers to the number of biological replicates and *n-Cells* refers to the number of individual cell measurements that passed all final filtering criteria across all biological replicates. The “FCS Mean” corresponds to the average nuclear concentration in nM measured for that condition. The “FCS SEM” is the standard error of the mean for the corresponding nuclear concentration measurement. Unless otherwise specified, the dosage is reported in the number of cas9 molecules per cell ($x10^7$) and in picomoles of Cas9 ribonucleoprotein complex per $10^5$ cells.

| **Cell Type** | **Time (hrs)** | **Dosage (Cas9 $x10^7$ per cell)** | **Dosage (pmol/$10^5$ cells)** | ***n* - BR** | ***n* - FCS** | **FCS Mean** | **FCS SEM** |
|---|---|---|---|---|---|---|---|
| HeLa | 12 | 7.5 | 12.5 | 2 | 28 | 22.1 | 1.10 |
| HeLa | 12 | 7.5 | 1.56 | 2 | 30 | 4.41 | 0.245 |
| HeLa | 24 | 15 | 25 | 2 | 33 | 38.6 | 2.58 |
| HeLa | 24 | 7.5 | 12.5 | 2 | 25 | 25.4 | 1.59 |
| HeLa | 24 | 7.5 (RNA only) | 12.5 (RNA only) | 2 | 26 | 19.4 | 0.843 |
| HeLa | 24 | 3.8 | 6.25 | 3 | 37 | 8.11 | 0.650 |
| HeLa | 24 | 1.9 | 3.125 | 2 | 28 | 3.98 | 0.243 |
| HeLa | 24 | 0.94 | 1.56 | 2 | 37 | 4.22 | 0.239 |
| HeLa | 24 | 0.24 | 0.4 | 2 | 34 | 2.94 | 0.287 |
| HeLa | 36 | 7.5 | 12.5 | 2 | 40 | 25.0 | 1.61 |

| | | | | | | | |
|---|---|---|---|---|---|---|---|
| HeLa | 36 | 0.94 | 1.56 | 2 | 42 | 4.12 | 0.191 |
| U2OS | 24 | 7.5 | 12.5 | 2 | 26 | 21.4 | 1.96 |
| U2OS | 24 | 0.94 | 1.56 | 2 | 21 | 6.86 | 2.10 |
| HEK293T | 24 | 7.5 | 12.5 | 2 | 20 | 30.0 | 1.98 |
| HEK293T | 24 | 0.94 | 1.56 | 2 | 20 | 6.28 | 0.521 |

***Supplementary Table 3: Antibodies used for Western Blots and Immunofluorescence microscopy***

Antibodies Used:

WB= Western Blot; IF = immunofluorescence; mAb = monoclonal Antibody; pAb = polyclonal antibody

Primary Antibodies:

| Antibody target/strain: | Source/Isotype: | Dilution(s) used: | Figures: | Company and Catalog # |
|---|---|---|---|---|
| CRISPR-Cas9 | Rabbit pAb | 1:1000 (WB), 1:100 (IF) | Fig. 4B (IF) | Diagenode (#C15310258) |
| FLAG (DDDDK sequence) | Rabbit Mab (recombinant) | 1:250 (IF), 1:2500 (WB) | Fig. 4C (IF), Fig. S6D (WB), Fig. S9A (IF) | Abcam (#ab205606) |
| HIV1 p24 | Mouse Mab | 1:250 (IF), 1:1000 (WB) | Fig. S6D (WB), Fig. S9A (IF) | Invitrogen (#MA1-71515) |

Secondary Antibodies:

| Antibody: | Dilution(s) used: | Figures: | Company and Catalog # |
|---|---|---|---|
| IRDye® 680RD Goat anti- | 1:2500 (WB) | Fig. S6D | Licor (Catalog # 926-68070) |

| | | | |
|---|---|---|---|
| Mouse IgG Secondary Antibody | | | |
| IRDye® 800CW Goat anti-Rabbit IgG Secondary Antibody | 1:2500 (WB) | Fig. S6D | Licor (Catalog # 926-32211) |
| Goat anti-Rabbit IgG (H+L) Highly Cross-Adsorbed Secondary Antibody, Alexa Fluor™ Plus 405 | 1:1000 (IF) | Fig. 4B | Invitrogen (Catalog # A48254) |
| Alexa Fluor™ 594 Tyramide SuperBoost™ Kit, goat anti-rabbit IgG | Used in accordance with manufacturer's instructions. | Fig. 4D, Fig. S9A | ThermoFisher (Catalog # B40944) |
| Alexa Fluor™ 488 Tyramide SuperBoost™ Kit, goat anti-mouse IgG | Used in accordance with manufacturer's instructions. | Fig. S9A | ThermoFisher (Catalog # B40941) |

| Counterstain: | Cellular Compartment: | Dilution(s) used: | Figures: | Company and Catalog # |
|---|---|---|---|---|
| SYTOX™ Deep Red Nucleic Acid Stain | Nucleus | 1:2000 (IF) | Fig. 4B,C | Invitrogen (#S11380) |
